# Lysosomes mediate the mitochondrial UPR via mTORC1-dependent ATF4 phosphorylation

**DOI:** 10.1101/2022.11.14.516427

**Authors:** Terytty Yang Li, Qi Wang, Arwen W. Gao, Xiaoxu Li, Adrienne Mottis, Minho Shong, Johan Auwerx

**Affiliations:** Laboratory of Integrative Systems Physiology, Interfaculty Institute of Bioengineering, École Polytechnique Fédérale de Lausanne, Lausanne, Switzerland; Division of Endocrinology and Metabolism, Department of Internal Medicine, Chungnam National University College of Medicine, Daejeon, Korea; Present address: Laboratory Genetic Metabolic Diseases, Amsterdam Gastroenterology, Endocrinology, and Metabolism, Amsterdam Cardiovascular Sciences, Amsterdam UMC, University of Amsterdam, Meibergdreef 9, 1105 AZ Amsterdam, The Netherlands

## Abstract

Lysosomes are central platforms for not only the degradation of macromolecules but also the integration of multiple signaling pathways. However, whether and how lysosomes mediate the mitochondrial stress response (MSR) remain largely unknown. Here, we demonstrate that lysosomal acidification via the vacuolar H^+^-ATPase (v-ATPase) is essential for the transcriptional activation of the mitochondrial unfolded protein response (UPR^mt^). Mitochondrial stress stimulates v-ATPase-mediated lysosomal activation of the mechanistic target of rapamycin complex 1 (mTORC1), which then directly phosphorylates the MSR transcription factor, activating transcription factor 4 (ATF4). Disruption of mTORC1-dependent ATF4 phosphorylation blocks the UPR^mt^, but not other similar stress responses, such as the UPR^ER^. Finally, ATF4 phosphorylation downstream of the v-ATPase/mTORC1 signaling is indispensable for sustaining mitochondrial redox homeostasis and protecting cells from reactive oxygen species (ROS)-associated cell death upon mitochondrial stress. Thus, v-ATPase/mTORC1-mediated ATF4 phosphorylation via lysosomes links mitochondrial stress to UPR^mt^ activation and mitochondrial function resilience.

---

Long known as the degradative end points for intra- and extra-cellular cargos, lysosomes have emerged as signaling centers that play important roles in nutrient sensing, cell growth, energy metabolism, immune response and aging^1–4^. Accordingly, dysfunction of lysosomes has been associated with a variety of diseases including lysosomal storage disorders, cancer, diseases of the immune system and neurodegenerative disorders^5–7^. To maintain energy homeostasis and protein quality control, lysosomes also constantly communicate with other cellular organelles, such as the mitochondria^8,9^. For example, severe mitochondrial dysfunction triggers mitophagy^10^, which results in the degradation of impaired mitochondria by the lysosomes; whereas changes in lysosomal pH or signaling may in turn modulate mitochondrial function and regulate longevity in different organisms^11,12^. However, whether and how the lysosomes mediate the communication from stressed mitochondria to the nucleus are still poorly understood.

The mitochondrial unfolded protein response (UPR^mt^), a branch of the mitochondrial stress response (MSR), is an adaptive transcriptional response that helps to resolve proteostatic toxicity triggered by diverse mitochondrial stresses^13–15^. Although first discovered in mammalian cells^14^, the regulatory mechanisms of UPR^mt^ have been particularly well-studied in the nematode *Caenorhabditis elegans*. In *C. elegans*, a panel of transcription factors/co-factors, histone methyltransferases, demethylases, acetyltransferases and deacetylase cooperate with the master UPR^mt^ transcription factor, activated transcription factor-1 (ATFS-1), to mediate the UPR^mt^ upon mitochondrial perturbations^14,16–19^. Nevertheless, how the mitochondrial stress signal is relayed through the cytosol and sensed by these UPR^mt^ regulators is largely unclear. In mammalian cells, mitochondrial stress triggers the integrated stress response (ISR)^20,21^, in which phosphorylation of the eukaryotic translation initiation factor 2α (EIF2α) results in the translation of several transcription factors including activating transcription factor 4 (ATF4), activating transcription factor 5 (ATF5) and C/EBP homologous protein (CHOP) to coordinate a gene expression program considered as the functional equivalent of the UPR^mt 13,14,22^. In a parallel study conducted in *C. elegans*, we revealed that increased ATFS-1 translation, mediated by the v-ATPase/TORC1 and lysosomes, contributes to the cytosolic relay of mitochondrial stress to direct UPR^mt^ activation through an EIF-2α phosphorylation-independent mechanism^23^. However, whether the roles of lysosomes and v-ATPase/TORC1 in UPR^mt^ regulation are evolutionally conserved remains elusive. Furthermore, how mammalian cells distinguish stress signals from different origins, such as the mitochondrion and ER, to concordantly activate the ATF4-mediated “integrated stress response” is still unknown.

## Suppression of lysosomal acidification inhibits UPR^mt^ in mammalian cells

The vacuolar H^+^-ATPase (v-ATPase) is a highly conserved large complex proton pump which locates at the lysosomal surface and is essential for the acidification of lysosomes^24,25^. In addition to its role as a proton pump, v-ATPase has also been shown to be crucial for the integration of multiple signaling pathways, including mechanistic target of rapamycin complex 1 (mTORC1)^1,26^, adenosine monophosphate-activated protein kinase (AMPK)^27^, as well as Janus kinase 2 (JAK2)-signal transducer and activator of transcription-3 (STAT3) signaling^28^. We first questioned if the role of v-ATPase in UPR^mt^ is functionally conserved in mammalian cells. Doxycycline (Dox)^17,29^, an antibiotic that inhibits mitochondrial ribosome translation, activated the MSR and increased the expression of many UPR^mt^ transcripts (e.g., *HSPA9*, *HSPD1* and *ASNS*) in human embryonic kidney (HEK) 293T cells (Fig. 1a). This response was suppressed by the knockdown of *ATP6V0C* and *ATP6V0D1* (Fig. 1a and Extended Data Fig. 1a), two core subunits of the v-ATPase complex^24,30^. Similarly, inhibition of v-ATPase activity by two small-molecule inhibitors, Bafilomycin A1 (BafA1) and Concanamycin A (ConA)^31,32^, strongly attenuated Dox-induced expression of typical UPR^mt^ genes in mouse embryonic fibroblasts (MEFs) (Fig. 1b and Extended Data Fig. 1b). Among these approaches to suppress UPR^mt^ activation, ConA treatment in MEFs was most efficacious (Fig. 1b). Strikingly, the half maximal inhibitory concentration (IC_50_) of ConA on inhibiting Dox-induced UPR^mt^ is below 1.5 nM in MEFs, while that of BafA1 is around 50 nM (Fig. 1c), in line with a more prominent effect of ConA in suppressing v-ATPase activity *in vitro*^32^.

**Fig. 1.**
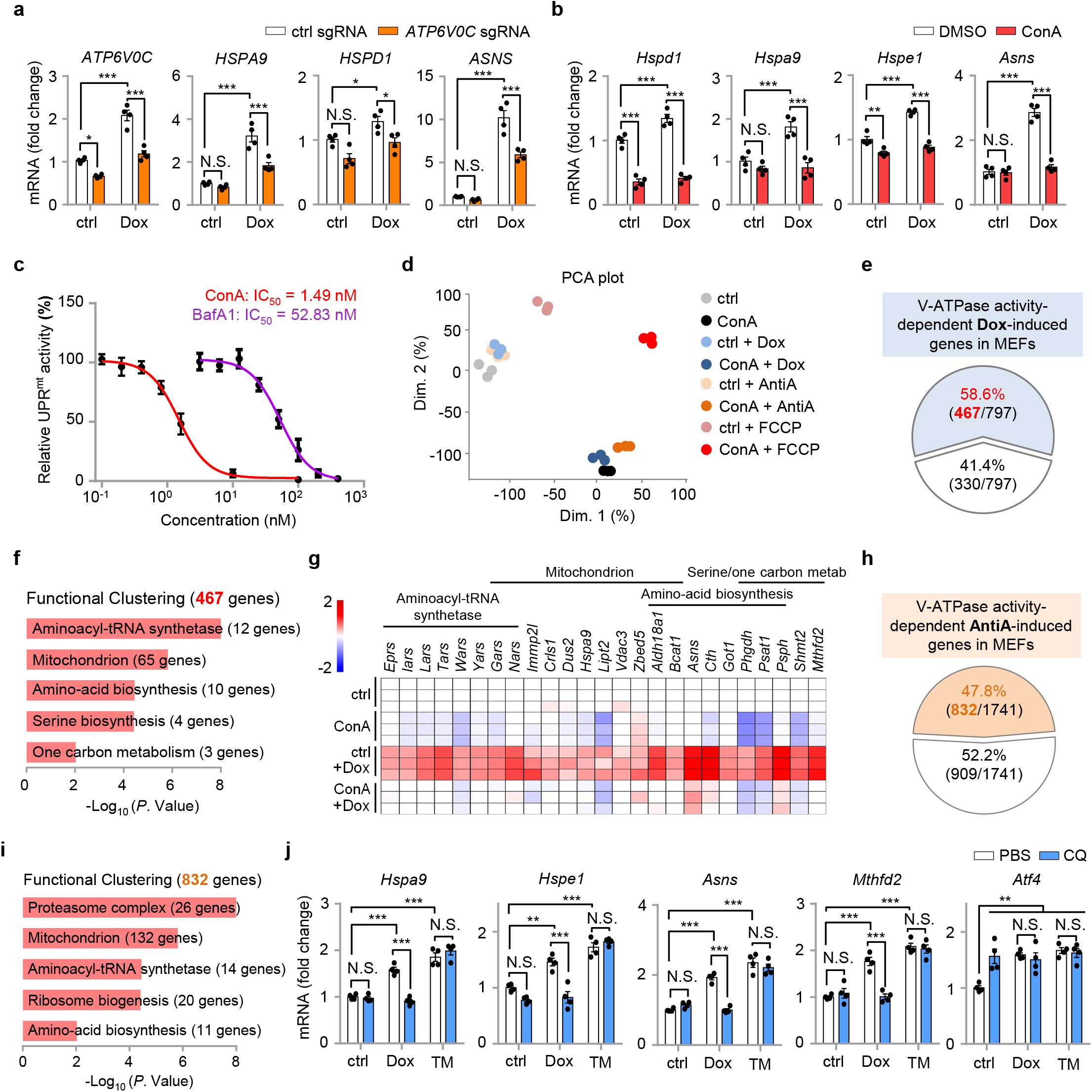
Suppression of lysosomal acidification inhibits UPR^mt^ activation in mammalian cells. **a**, qRT-PCR results (*n* = 4 biologically independent samples) of HEK293T cells expressing control (ctrl) or *ATP6V0C* sgRNA, and treated with or without Doxycycline (Dox) (30 μg/ml) for 24 h. **b**, qRT-PCR results (*n* = 4 biologically independent samples) of MEFs pretreated with DMSO control or v-ATPase inhibitor Concanamycin A (ConA) (200 nM) for 1 h, and then co-treated with or without Dox (30 μg/ml) for 24 h. **c**, The dose-response curve and IC_50_ of ConA (red) and BafA1 (purple) on inhibiting Dox-induced UPR^mt^ activation in MEFs. The relative UPR^mt^ activity was normalized to the mRNA induction level of *Hspa9* (*n* = 4 biologically independent samples) in response to Dox (30 μg/ml, 24 h), and with ConA or BafA1 co-treatment. **d**, Principal Component Analysis (PCA) of the RNA-seq profiles of MEFs treated with Dox, Antimycin A (AntiA), FCCP, and/or ConA. **e**, Diagram of the UPR^mt^ genes that are dependent (red) on v-ATPase activity for induction upon Dox treatment, according to the RNA-seq dataset. **f**, Functional clustering of the 467 v-ATPase activity-dependent genes as indicated in (**e**). **g**, Heat-map of the relative expression levels of representative v-ATPase activity-dependent UPR^mt^ genes in MEFs treated with Dox and/or ConA in log_2_ fold-change, based on the RNA-seq dataset. See Supplementary Table 1 for detailed gene expression changes. **h**, Diagram of the UPR^mt^ genes that are dependent (orange) on v-ATPase activity for induction upon AntiA treatment. **i**, Functional clustering of the 832 v-ATPase activity-dependent genes as indicated in (**h**). **j**, qRT-PCR results (*n* = 4 biologically independent samples) of MEFs pretreated with PBS control or 50 μM Chloroquine (CQ) for 1 h, and then co-treated with or without Dox (30 μg/ml) or Tunicamycin (TM, 1.5 μg/ml) for 24 h. Error bars denote S.E.M. Statistical analysis was performed by ANOVA followed by Tukey post-hoc test (\**P* < 0.05; \*\**P* < 0.01; \*\*\**P* < 0.001; N.S., not significant).

To systematically evaluate the impact of v-ATPase inhibition on the transcriptional activation of the MSR, we performed RNA sequencing (RNA-seq) on total RNA isolated from MEFs treated with DMSO control or ConA for 24 h, in the absence or presence of three mechanistically different mitochondrial stressors: a mitochondrial translation inhibitor, Dox^17,29^; a mitochondrial complex III inhibitor, Antimycin A (AntiA)^33,34^; and a mitochondrial oxidative phosphorylation uncoupler, carbonyl cyanide *p*-(trifluoromethoxy) phenylhydrazone (FCCP)^35^ (Fig. 1d). Dox up-regulated 797 transcripts (adjusted *P* value < 0.05), and 736 (92.3%) of them were also induced by AntiA or FCCP (Extended Data Fig. 1c and Supplementary Table 1). In contrast, FCCP led to the up-regulation of 4,182 transcripts, and only 1,364 (32.6%) of them were commonly shared with those induced by either Dox or AntiA (Extended Data Fig. 1c). As expected, the 443 genes up-regulated in response to all the three mitochondrial stress inducers were enriched for mitochondrial surveillance pathways such as “Aminoacyl-tRNA synthetase”, “Amino-acid biosynthesis” and “Mitochondrion” (Extended Data Fig. 1d), in line with previous studies^17,34,36^. Of note, in addition to “Mitochondrion”, the 2,818 transcripts that induced only by FCCP were also enriched for other organelles including “Golgi apparatus”, “Endosome” and “Endoplasmic reticulum (ER)” (Extended Data Fig. 1e), reminiscent of other metabolic impacts of FCCP or FCCP-like protonophore uncouplers through mechanisms irrelevant of mitochondrial membrane potential disruption^37,38^.

As expected, a more restricted number of transcripts was altered upon Dox stimulation in the presence of ConA (Extended Data Fig. 1f,g). Importantly, among the 797 Dox-induced transcripts, a majority (58.6%, 467 transcripts) of them was abrogated by ConA (Fig. 1e). These 467 genes, hereby defined as the “v-ATPase activity-dependent Dox-induced genes”, were enriched for MSR-related pathways (Fig. 1f), and include UPR^mt^ genes such as *Asns*, *Phgdh*, *Psat1*, *Psph*, *Shmt2* and *Mthfd2* (Fig. 1g). Likewise, around half (47.8%) of the AntiA-induced transcripts relied on v-ATPase activity for induction (Fig. 1h). Despite that only 30.4% of the FCCP-induced transcripts were abrogated by ConA (Extended Data Fig. 1h), the 832 AntiA-induced and 1,271 FCCP-induced transcripts that were dependent on v-ATPase activity were both enriched for mitochondrion-related pathways (Fig. 1i and Extended Data Fig. 1i). In contrast, the other 2,911 FCCP-induced transcripts independent on v-ATPase activity were highly enriched for other cellular organelles such as “Endoplasmic reticulum”, “Golgi apparatus”, “Lysosome” and “Endosome” (Extended Data Fig. 1j), suggesting that the v-ATPase inhibitor ConA probably attenuated the adaptive response specifically related to mitochondria, but not other cellular organelles such as the ER.

Consistently, ConA inhibited the induction of UPR^mt^ genes, including *Hspa9*, *Asns*, *Psph* and *Mthfd2*, upon exposure to mitochondrial stress inducers, AntiA and Oligomycin (Olig) (Extended Data Fig. 1k). Despite that the ER UPR (UPR^ER^) inducer Tunicamycin (TM) also up-regulated these MSR transcripts, as reported previously^39^, their induction upon TM treatment was surprisingly not affected by ConA (Extended Data Fig. 1k). Moreover, the TM-induced dramatic up-regulation of ER chaperone *Grp78/Bip*^40^, a key event of the UPR^ER 41,42^, was also not affected by ConA (Extended Data Fig. 1k). As an alternative approach to inhibit lysosome acidification and mimic v-ATPase loss-of-function, disruption of lysosomal pH gradient by chloroquine (CQ)^43^, also abrogated Dox-induced UPR^mt^ activation, but not TM-induced stress response (Fig. 1j). Interestingly, different from the effect of ConA and CQ, the mTOR inhibitors Rapamycin and Torin1^44^, robustly attenuated the induction of typical UPR^mt^ and UPR^ER^ genes in response to Dox or TM treatment^39^, and inhibited the expression of key transcription factors *Atf4*, *Atf5* and *Chop* (Extended Data Fig. 1l). Together, these results suggest that disruption of lysosomal acidification by inhibiting v-ATPase activity or CQ suppresses the UPR^mt^, but not other similar stress responses such as the UPR^ER^, while direct mTORC1 inhibition abrogates the transcriptional responses induced by both mitochondrial and ER stress inducers.

## Lysosomal inhibition increases ATF4 accumulation but also limits ATF4 binding to the promoters of UPR^mt^ genes

To investigate the mechanism on how lysosomes and v-ATPase regulate the UPR^mt^, we first checked the expression levels of the putative UPR^mt^ transcription factors (i.e., ATF4, ATF5 and CHOP) upon ConA treatment, with or without mitochondrial stress. Surprisingly, their mRNA levels were up-regulated even in cells with only ConA treatment (Extended Data Fig. 2a,b); the expression of multiple classical ATF4 targets, including *Chac1*, *Herpud1*, *Trib3* and *Slc7a11*, that are induced in response to ER stress^36,39^, were increased as well under this condition (Extended Data Fig. 2b). Similar patterns were also found for the mitophagy/autophagy transcripts (e.g., *Sqstm1*, *Binp3l*, *Pink1*) (Extended Data Fig. 2b). At protein level, Dox mildly increased EIF2α phosphorylation and ATF4 expression^36^ (Fig. 2a). Interestingly, more ATF4 protein was detected in ConA/Dox co-treated MEFs at all time points, as compared to Dox-only conditions (Fig. 2a). Meanwhile, the expression of ATF5 and CHOP was either not affected or only slightly induced by Dox or ConA (Fig. 2a). Moreover, ConA alone increased ATF4 protein expression in a time-dependent manner (Fig. 2b). In contrast, complete inhibition of mTORC1 activity by Torin1 led to depletion of ATF4 protein (Fig. 2b), in line with its mRNA changes and previous studies^45,46^ (Extended Data Fig. 1l).

**Fig. 2.**
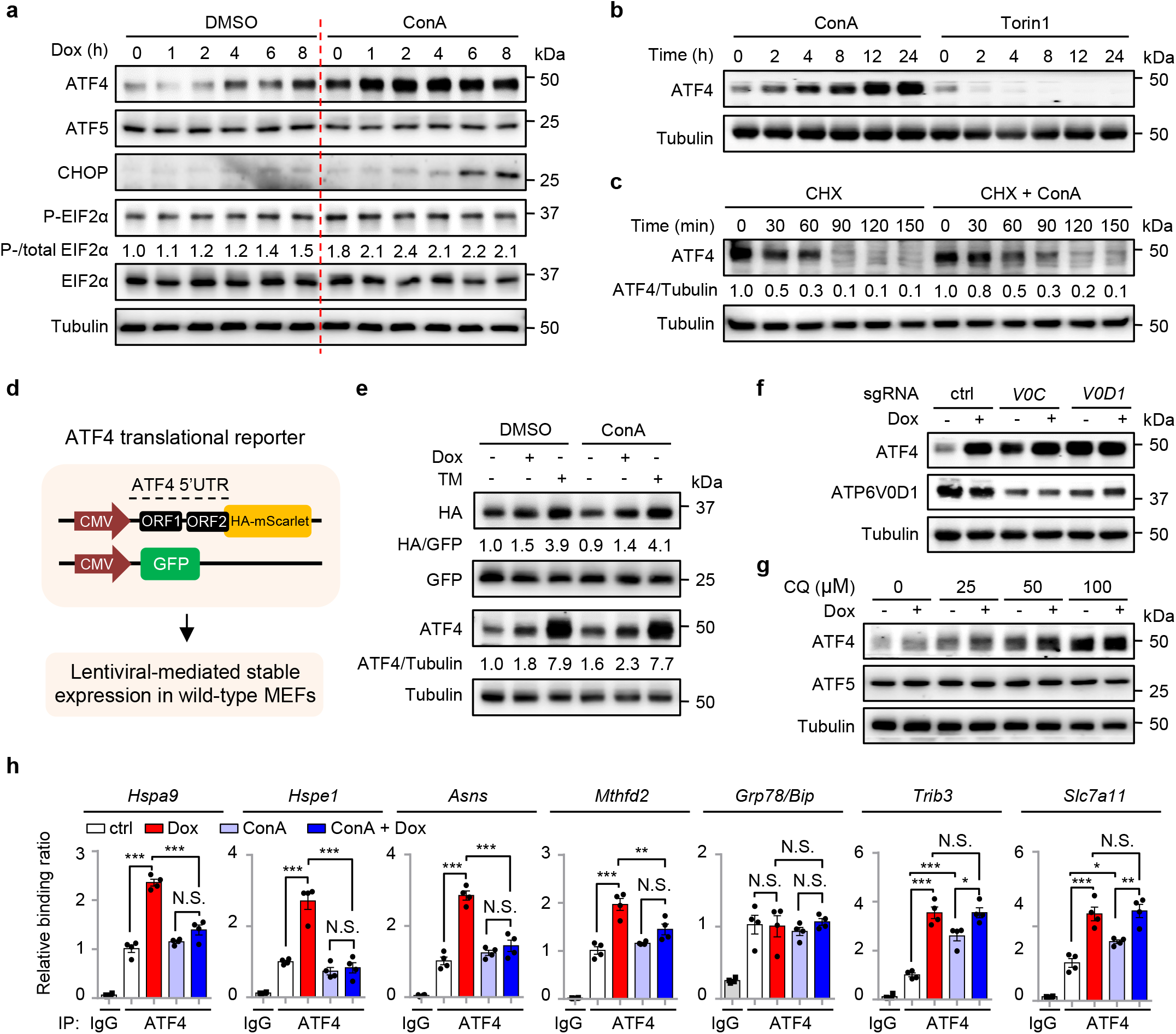
Lysosomal inhibition increases ATF4 accumulation but also limits ATF4 binding to the promoters of UPR^mt^ genes. **a**, Western blots showing time-dependent changes of proteins in MEFs pretreated with DMSO control or ConA (200 nM) for 1 h, and then co-treated with Dox (30 μg/ml) for 0-8 h. All ConA-treated conditions were thus treated with ConA for a total time of 9 h. **b**, Western blots of MEFs treated with ConA (200 nM) or Torin1 (250 nM) for 0-24 h. **c**, Western blots of MEFs treated with cycloheximide (CHX), in the absence or presence of ConA (200 nM) for 0-150 min. **d**, Schematic diagram of the ATF4 translational reporter, comprising the upstream open reading frames (uORF1 and uORF2) of the ATF4 5′ untranslated region (5′UTR) followed by HA-mScarlet tag replacing the ATF4 coding sequence, built on a lentiviral expression vector. The GFP control is directly driven by the cytomegalovirus (CMV) promoter. **e**, Western blots of MEFs stably expressing the ATF4 translational reporter and GFP control treated with or without Dox (30 μg/ml) or Tunicamycin (TM, 1.5 μg/ml) for 3 h, in the presence of DMSO control or ConA (200 nM). **f**, Western blots of HEK293T cells expressing control (ctrl), *ATP6V0C* (*V0C*) or *ATP6V0D1* (*V0D1*) sgRNA, and treated with or without Dox (30 μg/ml) for 24 h. **g**, Western blots of MEFs pretreated with PBS control or 25-100 μM chloroquine (CQ) for 1 h, and then co-treated with or without Dox (30 μg/ml) for 24 h. **h**, ATF4 ChIP-qPCR analysis (*n* = 4 biologically independent samples) of the promoters of ATF4 targeted genes in MEFs pretreated with DMSO or ConA (200 nM) for 1 h, and then co-treated with or without Dox (30 μg/ml) for 3 h. Error bars denote S.E.M. Statistical analysis was performed by ANOVA followed by Tukey post-hoc test (\**P* < 0.05; \*\**P* < 0.01; \*\*\**P* < 0.001; N.S., not significant).

To understand how ConA leads to ATF4 protein accumulation, we then measured the protein stability of ATF4 when protein synthesis was blocked by a translation inhibitor, cycloheximide (CHX). The half-life of ATF4 is very short at basal state^45,47^, and was almost doubled (from 30 to 60 min) in the presence of ConA (Fig. 2c). Next, by using a ATF4 translation reporter^48^, which expresses a HA-tagged reporter protein under the strict control of ATF4 5’UTR (Fig. 2d), we found that ConA in general did not affect the translation activity of ATF4 (as determined by comparing the expression of the ATF4 5’UTR-driven HA-tagged reporter protein and the normal CMV5 promoter-driven GFP protein), in response to either Dox or TM treatment for 3 h (Fig. 2e). Meanwhile, the endogenous expression of ATF4 protein increased by 60% upon only ConA exposure (Fig. 2e). These results suggest that a basal mTORC1 activity still retains for normal ATF4 translation even with v-ATPase inhibition, and ConA increases ATF4 accumulation through a translation-independent mechanism.

Increased expression of ATF4 was also found in cells with *ATP6V0C* or *ATP6V0D1* knockdown, or CQ treatment (Fig. 2f,g). To elucidate how the accumulated ATF4 protein in ConA-treated MEFs failed to activate the UPR^mt^ genes upon mitochondrial stress (Fig.1g and Supplementary Table 1), we pulled-down the endogenous ATF4 in MEFs with or without ConA and/or Dox treatment, and performed a chromatin immunoprecipitation coupled with quantitative PCR (ChIP-qPCR). Dox strongly promoted the enrichment of ATF4 at the loci of multiple UPR^mt^ genes (e.g., *Hspa9*, *Hspe1*, *Asns*, *Mthfd2*), which was almost completely blocked by ConA (Fig. 2h). In contrast, enrichment of ATF4 at the loci of UPR^ER^-related genes was either unchanged (e.g., *Grp78/Bip*) or even increased (e.g., *Trib3*) upon ConA (Fig. 2h), in line with their changes at mRNA level (Extended Data Fig. 2b). Thus, inhibition of lysosomal acidification increases ATF4 accumulation but also limits ATF4 binding to the promoters of UPR^mt^ genes during mitochondrial stress.

## Mitochondrial stress induces a lysosomal activity-dependent mTORC1 activation at the lysosomal surface

Next, we checked whether mTORC1 signaling is activated upon mitochondrial stress, as revealed in our recent studies in human thyroid cancer cells and in *C. elegans*^23,49^. mTORC1 activity (as reflected by the phosphorylation of S6K, S6 and 4E-BP1) increased and peaked at 2-4 h of Dox treatment in MEFs, which was attenuated by the v-ATPase inhibitor ConA (Fig. 3a). A similar time-dependent activation pattern of mTORC1 was also observed in MEFs treated with AntiA (Fig. 3b), or with Olig treatment as reported eleswhere^50^. In contrast, the ER stress inducer TM gradually decreases S6K phosphorylation in MEFs (Fig. 3b), consistent with previous studies^39,51^. In line with the model that a mitochondrion-endosome-lysosome route that shuttles cargo from mitochondria to lysosomes is activated upon oxidative stress^52^, the mitochondrial dye MitoTracker^53^ strongly co-localized with early endosome vesicles (Rab5^+^), and partially with late endosome vesicles (Rab7^+^), but not with mature lysosomes (Lamp1^+^), 3 h after Dox treatment (Extended Data Fig. 3a-c). This phenomenon is likely caused by the breakdown of mitochondria in the more acidic late endosomes or lysosomes, which in turn resulted in the loss of the mitochondrial signal^52,54^. mTORC1 activation requires its dynamic recruitment to the lysosomal surface^26^. As expected, both Dox and AntiA promotes lysosomal, but not early or late endosomal, localization of mTORC1, which was furthermore suppressed by ConA (Fig. 3c and Extended Data Fig. 3d,e). Finally, increased mTORC1 activity, as reflected by the S6 phosphorylation, was also detected *in vivo* in the kidneys of wild-type C57BL/6J mice upon Dox administration (Fig. 3d), in accordance with the up-regulation of UPR^mt^ genes (Fig. 3d,e). These results suggest that mitochondrial stress induces a time-dependent activation of mTORC1 signaling, which furthermore relies on the intact function of v-ATPase and the lysosomes.

**Fig. 3.**
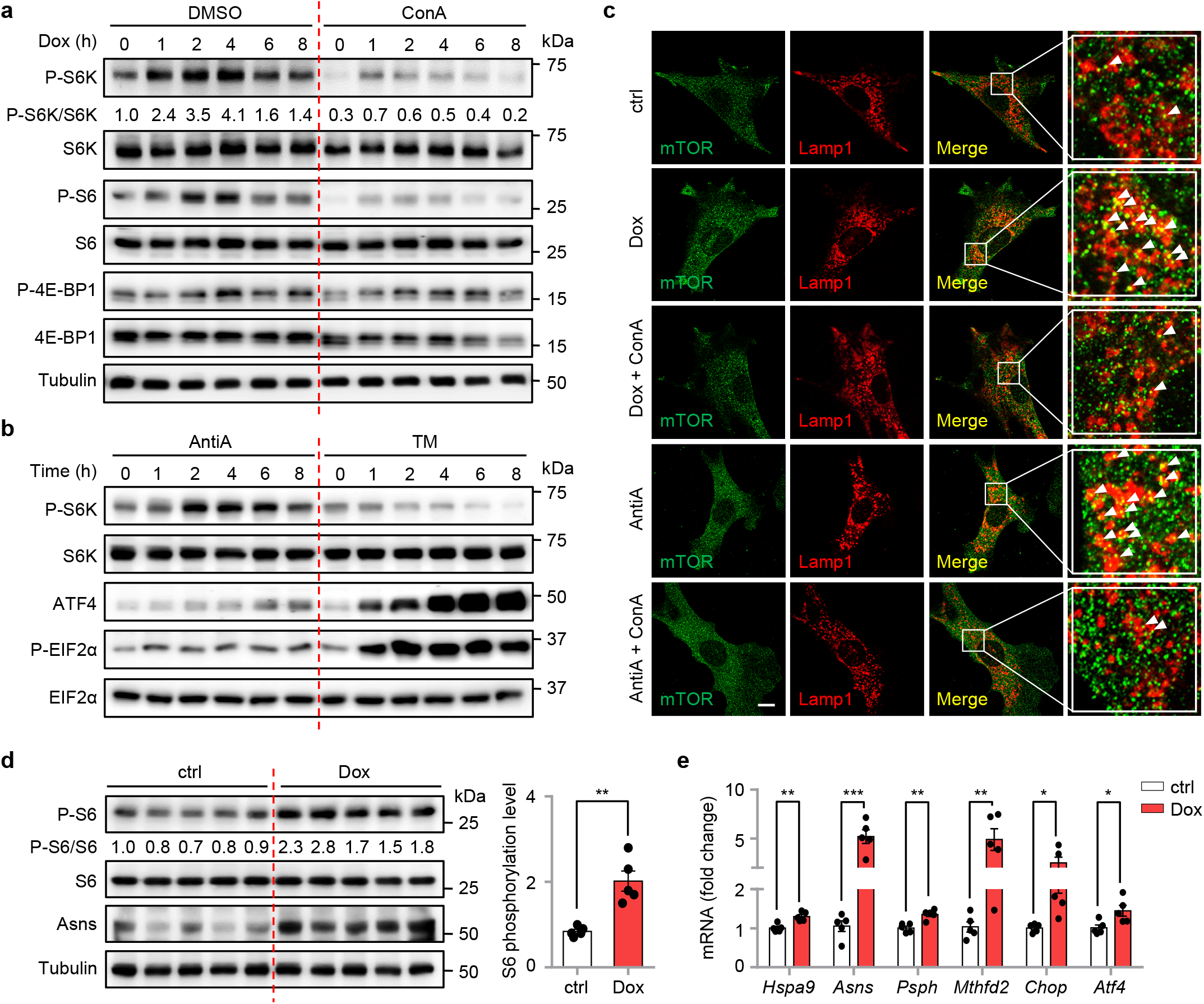
Mitochondrial stress induces v-ATPase-dependent mTORC1 activation at the lysosomal surface. **a**, Western blots showing time-dependent changes of proteins in MEFs pretreated with DMSO or ConA (200 nM) for 1 h, and then co-treated with or without Dox (30 μg/ml) for 0-8 h. **b**, Western blots showing time-dependent changes of proteins in MEFs treated with Antimycin A (AntiA, 2 μM) or Tunicamycin (TM, 1.5 μg/ml) for 0-8 h. **c**, ConA inhibits mitochondrial stress-induced lysosomal localization of mTOR in MEFs. MEFs were pretreated with DMSO control or ConA (200 nM) for 1 h, and then co-treated with or without Dox (30 μg/ml) for 3 h, cells were then fixed and co-stained with mTOR (green) and lysosome marker Lamp1 (red) antibodies. The arrows indicate the mTOR-lysosome co-localized puncta. Scale bar, 10 μm. **d**,**e**, Western blots (**d**) and qRT-PCR results (*n* = 5 mice for each group) (**e**) of kidney samples from 9-10 weeks-old male C57BL/6J mice treated with vehicle control (ctrl) or Dox (50 mg/kg) for 24 h. Error bars denote S.E.M. Statistical analysis was performed by two-tailed unpaired Student’s *t*-test (\**P* < 0.05; \*\**P* < 0.01; \*\*\**P* < 0.001; N.S., not significant).

## ATF4 is a direct phosphorylation target of mTOR in response to mitochondria stress

The fact that the accumulated ATF4 in ConA-treated MEFs failed to activate UPR^mt^ hints to the existence of certain post-translational modifications of ATF4, which are regulated in a v-ATPase/mTORC1-dependent fashion. In light of the kinase nature of mTOR, we questioned whether ATF4 is a direct phosphorylation target of mTORC1. Co-expression of ATF4 with the mTORC1 upstream activator Rheb increased the phosphorylation signal that was detected by a context-dependent (S*P) phosphorylation-specific antibody, which was inhibited by Torin1 (Fig. 4a). In contrast, no apparent phosphorylation signal was found when ATFS-1, the master transcription factor of UPR^mt^ in *C. elegans*^16^, was co-expressed with or without Rheb (Fig. 4a). Importantly, mTORC1-dependent phosphorylation of ATF4 was also detected in an *in vitro* kinase assay using either the mTORC1 immuno-precipitated from HEK293T cells or a recombinant kinase active mTOR protein purified from insect cells (Fig. 4b and Extended Data Fig. 4a). Mass spectrometric analysis revealed the existence of 5 serine/threonine (S/T) sites on ATF4 that can be phosphorylated by mTOR, and are sensitive to Torin1 treatment (Fig. 4c and Extended Data Fig. 4b). These sites are in general well-conserved across vertebrate species (Fig. 4d). It has been reported that mTORC1 substrates including S6K and 4E-BP1 harbor a canonical five amino acid TOR signaling (TOS) motif that is crucial for their regulation by mTORC1^55,56^. We discovered that a highly conserved TOS motif (-FDLDA-) also exists in the N-terminal of ATF4 protein (Fig. 4d), supporting that ATF4 is a *bona fide* evolutionally conserved phosphorylation target of mTOR. Separated or combined mutation of the 5 phosphorylation candidate sites to alanine revealed that ATF4 phosphorylation at Ser^166^ was specifically recognized by the context-dependent (S*P) phosphorylation antibody (referred hereafter as P-S166-ATF4 antibody), while P-T173-ATF4 was revealed by another context-dependent (ST*P) antibody (Fig. 4e). Moreover, Dox increased the endogenous P-S166-ATF4 and P-T173-ATF4 levels, which were furthermore abrogated by both ConA and Torin1 (Fig. 4f). Finally, increased phosphorylation of ATF4 at S166 and T173 was also detected upon exposure to the mitochondrial stress inducers such as AntiA and Olig (Fig. 4g), while ER stress inducer TM suppressed ATF4 phosphorylation, consistent with the changes of S6K phosphorylation (Fig. 4g). Thus, ATF4 is a direct phosphorylation target of mTORC1 downstream of multiple UPR^mt^ activators, but not the UPR^ER^ inducer TM.

**Fig. 4.**
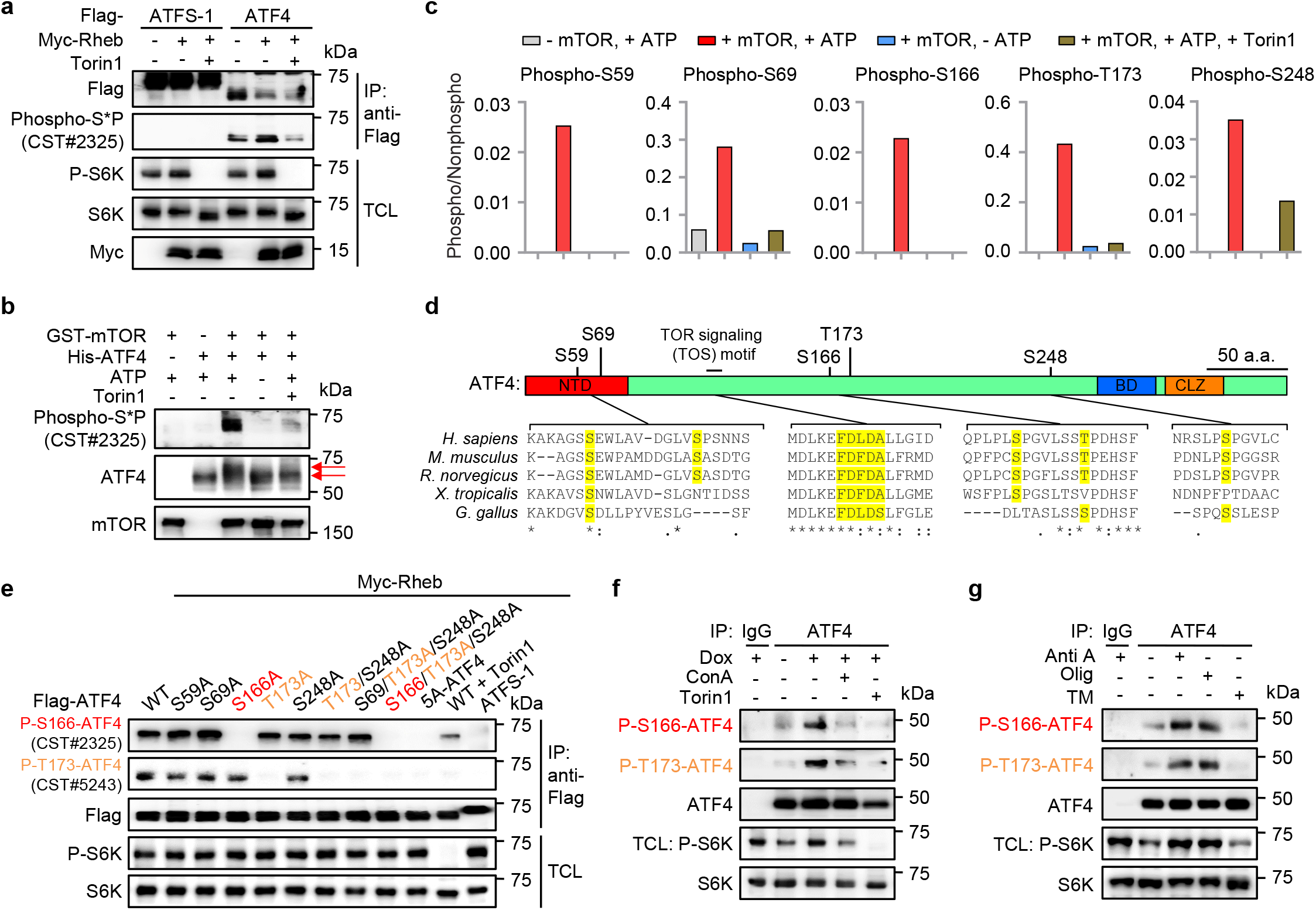
ATF4 is a direct phosphorylation target of mTOR in response to mitochondria stress. **a**, ATF4 phosphorylation recognized by a context-dependent (S*P) phosphorylation-specific antibody is increased by Rheb co-expression and inhibited by Torin1. HEK293T cells transfected with the indicated plasmids were immuno-precipitated (IP) with anti-Flag antibody and analyzed by western blots. When applicable, Torin1 (250 nM) were added 2 h before harvest. TCL, total cell lysate. **b**, mTOR directly phosphorylates ATF4 *in vitro*. *In vitro* kinase assay was performed with recombinant GST-tagged human mTOR purified from baculovirus-infected insect cells and recombinant His-tagged human ATF4 with or without Torin1 (250 nM). Arrows indicate the mobility shifts likely separating the hyperphosphorylated and nonphosphorylated ATF4. **c**, The ratios of the phosphorylated and nonphosphorylated peptides containing the phosphorylation sites of ATF4 from a kinase assay performed similar to (**b**), as determined by mass spectrometry. **d**, The identified mTOR-targeted phosphorylation sites on ATF4 with the vertebrate orthologs aligned below, with numbering according to the amino acid sequence of human ATF4 protein. NTD, N-terminal domain; BD, Basic domain; CLZ, C-leucine zipper. The highly conserved putative TOR signaling (TOS) motif was also highlighted. **e**, Validation of the two commercially available antibodies that specifically recognize ATF4 phosphorylation at Ser^166^ and Thr^173^, respectively. HEK293T cells transfected with the indicated plasmids were immuno-precipitated with anti-Flag antibody and analyzed by western blots. Torin1 (250 nM) were added 2 h before harvest. **f**, Increased ATF4 Ser^166^ and Thr^173^ phosphorylation upon Dox treatment, which was inhibited by ConA and Torin1. Wild-type MEFs were pretreated with DMSO, ConA (200 nM) or Torin1 (250 nM) for 1 h, and then co-treated with or without Dox (30 μg/ml) for 2 h, immuno-precipitated with anti-ATF4 antibody and analyzed by western blots. **g**, Increased ATF4 phosphorylation upon mitochondrial, but not ER stress inducers. Wild-type MEFs were with treated with Antimycin A (AntiA, 2 μM), Oligomycin (Olig, 2 μM), or Tunicamycin (TM, 1.5 μg/ml) for 2 h, immuno-precipitated with anti-ATF4 antibody and analyzed by western blots. Similar amount of immuno-precipitated ATF4 protein was loaded for different conditions to compare phosphorylation changes in (**f**) and (**g**).

## An essential role of ATF4 and its phosphorylation by mTORC1 in UPR^mt^ activation

Characterization of *Atf4*^−/−^ MEFs revealed an essential role of ATF4 in the induction of typical UPR^mt^ genes in response to Dox or AntiA treatment (Fig. 5a). Furthermore, certain UPR^mt^ genes (e.g., *Asns*, *Psph*, *Mthfd2*) rely on ATF4 for basal expression (Fig. 5a), in line with a previous study^57^. While the basal oxygen consumption rate (OCR) was remarkably decreased in *Atf4*^−/−^ MEFs as compared to that in WT cells, the maximum OCR after acute FCCP treatment was not affected (Fig. 5b), supporting a global metabolic reprogramming upon *ATF4* loss-of-function^36,57^. Whereas in the basal state, the mitochondrial network in *Atf4*^−/−^ MEFs is similar to that in WT cells (Fig. 5c), the mitochondrial network tended to be more disrupted in *Atf4*^−/−^ MEFs upon Dox administration, an effect that was even more pronounced after AntiA exposure, (Fig. 5c). The different impacts of Dox and AntiA on the mitochondrial network are also in line with a more potent effect of AntiA in regulating gene expression and UPR^mt^ activation (Fig. 5a and Extended Data Fig. 1c).

**Fig. 5.**
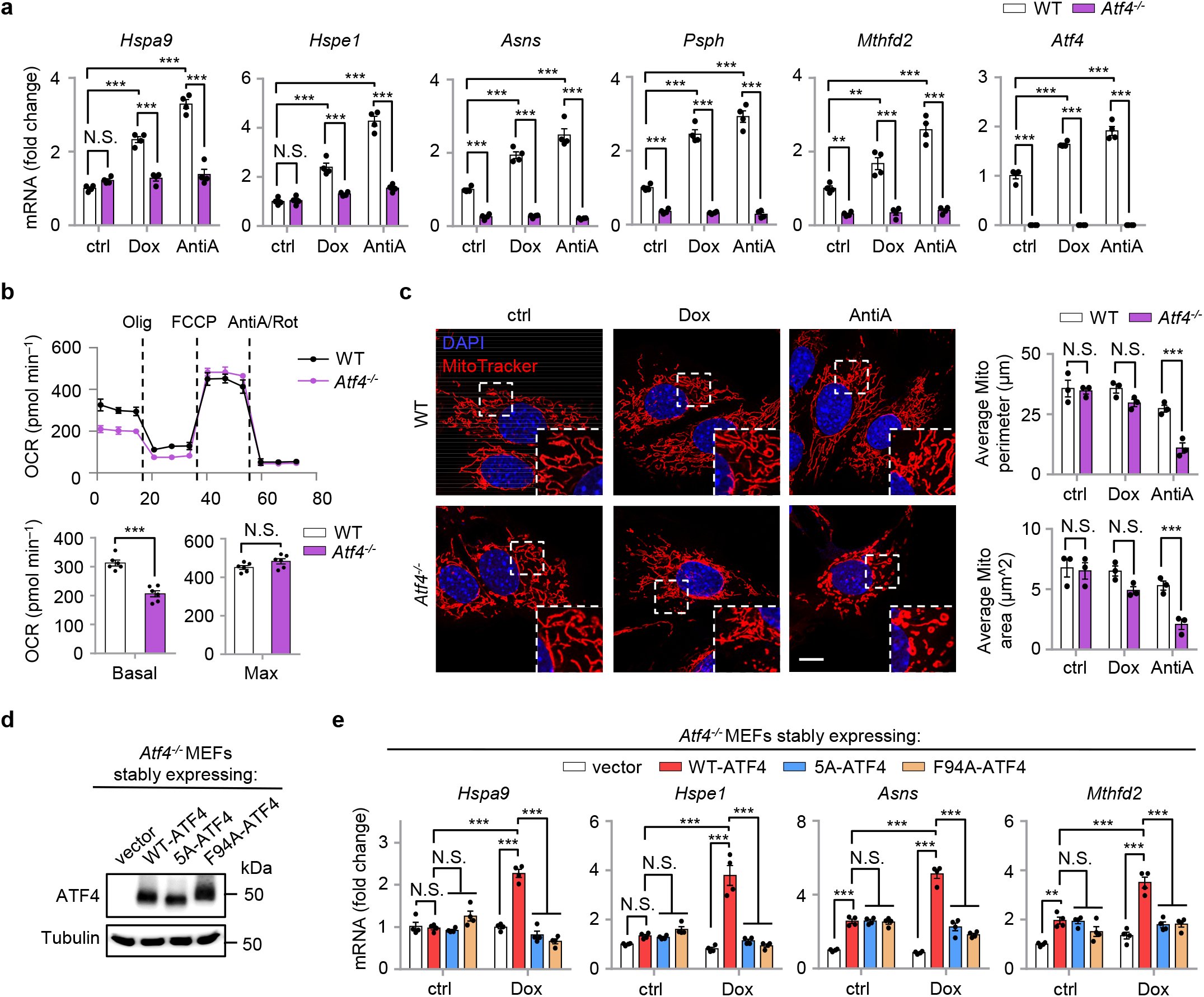
An essential role of ATF4 and its phosphorylation by mTORC1 in UPR^mt^ activation. **a**, qRT-PCR results (*n* = 4 biologically independent samples) of wild-type (WT) and *Atf4^−/−^* MEFs treated with or without Dox (30 μg/ml) or Antimycin A (AntiA, 2 μM) for 24 h. **b**, The oxygen consumption rate (OCR) of WT or *Atf4^−/−^* MEFs at basal or after sequential addition of Oligomycin (Olig), FCCP and AntiA/Rotenone. The basal and maximum OCR was statistically analyzed (*n* = 6 biologically independent samples). **c**, *Atf4* knockout leads to disrupted mitochondrial network upon mitochondrial stress. MitoTracker staining of WT or *Atf4^−/−^* MEFs treated with or without Dox (30 μg/ml) or AntiA (2 μM) for 24 h. The average mitochondrial network perimeter and area were analyzed by ImageJ with a Mito-Morphology macro (*n* = 3 independent experiments). Scale bar, 10 μm. **d**, Western blots of *Atf4^−/−^* MEFs stably expressing empty vector (vector), the wild-type ATF4 (WT-ATF4), the phospho-defective mutant (5A-ATF4), and an ATF4 mutant carrying a point mutation of the bulky phenylalanine residue 94 in the TOS motif to alanine (F94A-ATF4). **e**, qRT-PCR results (*n* = 4 biologically independent samples) of *Atf4^−/−^* MEFs stably expressing vector, wild-type, 5A or F94A forms of ATF4, treated with or without Dox (30 μg/ml) for 24 h. Error bars denote S.E.M. Statistical analysis was performed by two-tailed unpaired Student’s *t*-test in (**b**), or by ANOVA followed by Tukey post-hoc test in (**a**), (**c**) and (**e**) (\*\**P* < 0.01; \*\*\**P* < 0.001; N.S., not significant).

We then reconstituted the *Atf4*^−/−^ MEFs with either an empty vector control (vector), wild-type ATF4 (WT-ATF4), the ATF4 phosphorylation defective mutant (5A-ATF4, with all 5 serine/threonine phosphorylation sites mutated to alanine) or an ATF4 mutant carrying a point mutation of the bulky phenylalanine residue 94 (numbering for human ATF4) in the TOS motif to alanine (F94A-ATF4) (Fig. 5d). The phenylalanine to alanine mutation in the TOS motif of canonical mTOR substrates typically disrupts the functional impact of mTORC1 on its substrates, including on S6K and 4E-BP1^55,56^. As expected, in contrast to WT-ATF4, both 5A-ATF4 and F94A-ATF4 were unable to activate the UPR^mt^ in *Atf4*^−/−^ MEFs upon Dox treatment (Fig. 5e). Moreover, these ATF4 mutants failed to increase their binding to the promoters of UPR^mt^ genes in response to Dox (Extended Data Fig. 5a), in line with the results with ConA treatment (Fig. 2h). Furthermore, the TM-induced expression of the representative UPR^mt^/UPR^ER^ shared genes (i.e., *Hspa9*, *Hspe1*, *Asns* and *Mthfd2*) or the UPR^ER^-specific target *Grp78/Bip* was not affected in 5A-ATF4 or F94A-ATF4-rescued *Atf4*^−/−^ MEFs, relative to WT-ATF4-rescued MEFs (Extended Data Fig. 5b). Meanwhile, the expression of *Hspa9* and *Hspe1* still significantly increased upon TM treatment, even in *Atf4^−/−^* MEFs rescued with the empty control vector (Extended Data Fig. 5b), confirming the involvement of other transcription factors (e.g., ATF6 and XBP1s) in ER stress response^41,42,57^. Thus, both ATF4 and its phosphorylation by mTORC1 are essential for mitochondrial stress-induced UPR^mt^ activation.

## mTORC1-mediated ATF4 phosphorylation sustains mitochondrial homeostasis and protects cells from ROS-associated cell death upon mitochondrial stress

We then assessed the role of mTORC1-mediated ATF4 phosphorylation in mitochondrial homeostasis and function. In line with the findings in *Atf4*^−/−^ MEFs (Fig. 5c), healthy and well-connected mitochondrial network still exists in *Atf4*^−/−^ MEFs reconstituted with WT-ATF4, the 5A-ATF4 or F94A-ATF4 at basal state (Fig. 6a). A trend towards disrupted mitochondrial network upon Dox treatment was seen in 5A-ATF4- or F94A-ATF4-rescued *Atf4*^−/−^ MEFs, but not in those rescued with WT-ATF4; this tendency became more pronounced after the cells were challenged with AntiA (Fig. 6a). Consistently, reduced OCR were found in 5A-ATF4 or F94A-ATF4-rescued *Atf4*^−/−^ MEFs, compared to that in MEFs rescued with WT-ATF4 after AntiA exposure (Fig. 6b). By tracking the mitochondrial reactive oxygen species (ROS) level with MitoSOX^58^, remarkably higher percentages of MitoSOX positive cells were detected in *Atf4*^−/−^ MEFs expressing 5A-ATF4 and F94A-ATF4 after AntiA treatment, compared to that in cells expressing WT-ATF4 (Fig. 6c). Finally, the *Atf4^−/−^* MEFs reconstituted with 5A-ATF4 or F94A-ATF4 were more prone to AntiA-induced cell death, which was rescued with the supplement of the antioxidant, β-mercaptoethanol (β-ME) (Fig. 6d). Of note, in contrast to the *Atf4^−/−^* MEFs which require the addition of both the non-essential amino acids and certain antioxidants (e.g., β-ME) in the culture media to maintain survival^57^, *Atf4*^−/−^ MEFs rescued with 5A-ATF4 or F94A-ATF4 grow as well as those rescued with WT-ATF4 even without these supplements at unstressed condition (Fig. 6d). We have also noticed that *Atf4*^−/−^ MEFs rescued with F94A-ATF4 demonstrated more severe defects in mitochondrial function upon mitochondrial stress (Fig. 6a-c), as compared to that in cells rescued with 5A-ATF4, suggesting that there may exist other phosphorylation sites on ATF4 targeted by mTORC1 in addition to the 5 Serine/Threonine sites that we have identified. Together, these results indicate that while disruption of mTORC1-mediated ATF4 phosphorylation does not affect the basal function of ATF4 in maintaining cell growth, ATF4 phosphorylation downstream of v-ATPase/mTORC1 signaling plays a determining role in sustaining mitochondrial homeostasis and promoting survival from ROS-associated cell death in response to mitochondrial stress.

**Fig. 6.**
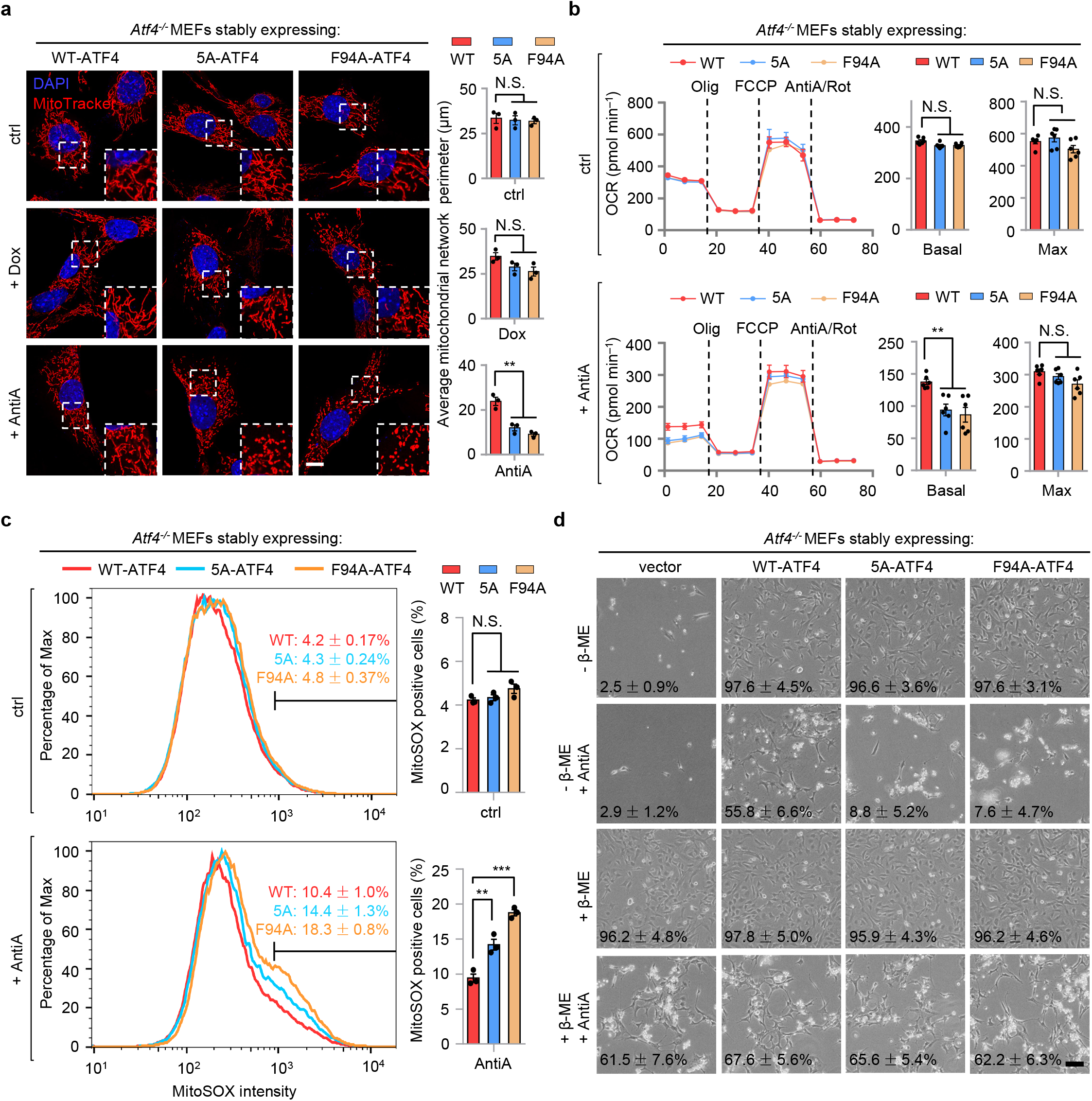
mTORC1-mediated ATF4 phosphorylation sustains mitochondrial homeostasis and protects cells from ROS-associated cell death upon mitochondrial stress. **a**, MitoTracker staining of *Atf4^−/−^* MEFs stably expressing wild-type (WT), 5A or F94A forms of ATF4, with or without Dox (30 μg/ml) or AntiA (2 μM) treatment for 24 h. The average mitochondrial network perimeter and area were analyzed by ImageJ with a Mito-Morphology macro (*n* = 3 independent experiments). Scale bar, 10 μm. **b**, The oxygen consumption rate (OCR) of *Atf4^−/−^* MEFs stably expressing WT, 5A or F94A forms of ATF4, after DMSO control (ctrl) or AntiA (2 μM) treatment for 24 h. The basal and maximum OCR was analyzed (*n* = 6 biologically independent samples). **c**, Flow cytometry analysis of the mitochondrial superoxide (MitoSOX) intensity of *Atf4^−/−^* MEFs stably expressing WT, 5A or F94A forms of ATF4, after DMSO control or AntiA (2 μM) exposure for 48 h. The percentages of MitoSOX-positive cells were analyzed (*n* = 3 independent experiments). **d**, Representative bright field photographs of *Atf4^−/−^* MEFs stably expressing empty vector, WT, 5A or F94A forms of ATF4, grown with or without the antioxidant β-mercaptoethanol (β-ME) or AntiA (2 μM) for 96 h. Mean percentages (± S.E.M) of the survival ratio of cells are indicated (*n* = 3 independent experiments). Error bars denote S.E.M. Statistical analysis was performed by ANOVA followed by Tukey post-hoc test (\*\**P* < 0.01; \*\*\**P* < 0.001; N.S., not significant).

## Discussion

The activity of mTORC1 has been shown to be required for increased ATF4 expression downstream of growth signals and during mitochondrial myopathy^39,46,59^. Full inhibition of mTORC1 furthermore induces the rapid loss of ATF4 at both mRNA and protein levels^45,46^, which likely explains why previous phosphoproteomic screens for mTORC1 substrates did not manage to identify ATF4 as one of the mTORC1 phosphorylation target^60,61^, since much less ATF4 will be detected after Torin1/Rapamycin exposure. This rapid loss of ATF4 expression hence blurs the correlation of phosphorylated and nonphosphorylated ATF4 peptides before and after mTORC1 inhibition with mTORC1 activity in these two screens^45,46^. Thus, the mechanism on how mTORC1 regulates ATF4 function, especially during mitochondrial stress, is still not fully understood. Moreover, due to the fact that the UPR^mt^ is often considered as part of the ISR downstream of the EIF2α phosphorylation event in mammalian systems^41,42^, it has to be determined how cells distinguish stress signals from different origins, i.e., mitochondrion and ER, to accurately activate the distinct UPR^mt^ and UPR^ER^ programs respectively.

Here, we reveal that mitochondrial stress and ER stress activate mechanistically different pathways, involving the lysosomes, v-ATPase/mTORC1, ATF4 and/or ribosomes, to concordantly activate the UPR^mt^ and the UPR^ER^ (Fig. 7). We found that in response to mitochondrial stress, mTORC1 is activated at the lysosomal surface via the v-ATPase, whereas EIF2α phosphorylation is only mildly increased, leading to a moderate increase in ATF4 translation (Figs. 2e, 3b). Meanwhile, activated mTORC1 directly phosphorylates ATF4, leading to increased ATF4 binding to the promoters of UPR^mt^ genes and the activation of UPR^mt^. In contrast, upon ER stress, mTORC1 activity is gradually suppressed but EIF2α phosphorylation is robustly increased, leading to a robust increase in ATF4 translation, and the subsequent activation of the UPR^ER^. Disruption of lysosomal acidification by CQ or the v-ATPase inhibitor ConA hence specifically suppressed the activation of the UPR^mt^ but not of the UPR^ER^. Additionally, in *C. elegans*, either v-ATPase or TORC1 suppression merely abrogated the UPR^mt^, but not UPR^ER^ or cytosolic UPR (UPR^CYT^)^23^. Thus, v-ATPase acts as an evolutionally conserved node relaying the stress signal specifically from mitochondria, but not ER, to the nuclear transcriptional adaptive response.

**Fig. 7.**
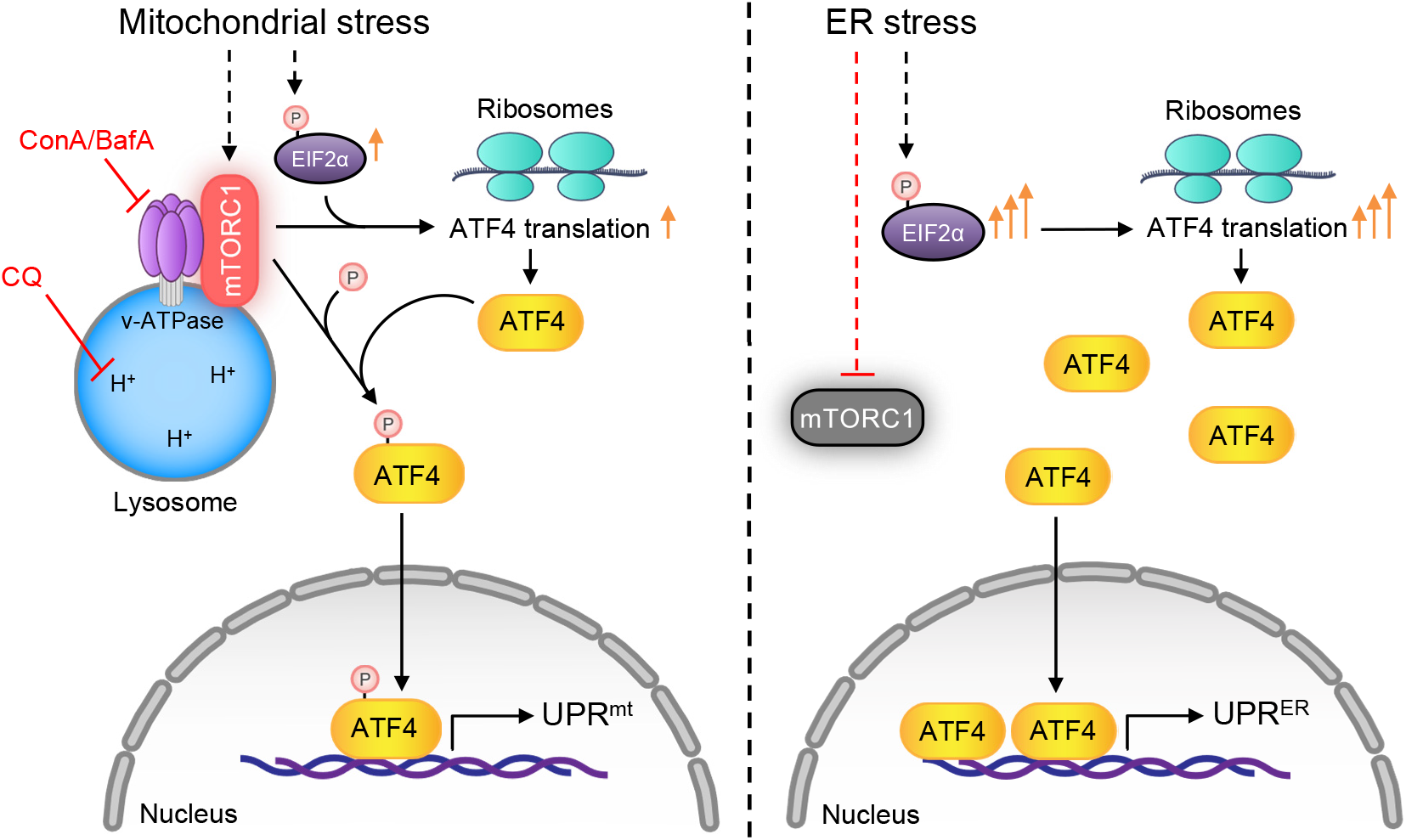
Models of how mitochondrial stress and ER stress activate mechanistically different pathways, involving the lysosomes, v-ATPase/mTORC1, ATF4 and/or ribosomes, to concordantly activate the UPR^mt^ and the UPR^ER^. Left: in response to mitochondrial stress, mTORC1 is activated at the lysosomal surface and EIF2α phosphorylation is mildly increased, leading to moderate increase in ATF4 translation. Meanwhile, activated mTORC1 directly phosphorylates ATF4, leading to increased ATF4 binding to the promoters of mitochondrial UPR (UPR^mt^) genes and UPR^mt^ activation. Right: in response to ER stress, mTORC1 activity is suppressed but the EIF2α phosphorylation is dramatically increased, leading to strongly increased translation and nuclear accumulation of ATF4, and subsequent activation of ER UPR (UPR^ER^). CQ, chloroquine. ConA/BafA1, Concanamycin A/Bafilomycin A1.

How exactly mitochondrial stress leads to the activation of the v-ATPase-mTORC1-ATF4 signaling remains an important direction for future work. One possibility is that the unfolded proteins/peptides accumulated during mitochondrial stress may somehow be transported from the mitochondria to the lysosomes, and get further digested to amino acids inside the lysosomes, which then activates mTORC1 via the v-ATPase^26,62,63^. Of note, mitochondrial stress, together with the unfolded mitochondrial-derived proteins/peptides, likely represents a unique intrinsic signal for mTORC1 activation by lysosome-derived amino acids^64^, which apparently differs from what is observed in response to growth factors or exogenous amino acids^64,65^. Accordingly, more mitochondrial proteins/peptides were detected in Rab5-positive endosomes after mitochondrial perturbatins^66^. Moreover, a Rab5-mediated mitochondrion-endosome-lysosome pathway was activated during mitochondrial redox stress, and functions in mitochondrial quality control independent of the mitophagy process^54,67^.

Collectively, our findings identified mammalian ATF4 as a direct phosphorylation target of mTORC1, and revealed a pivotal role of lysosomes and the v-ATPase/mTORC1 complex in mediating stress signal sensing and transduction from mitochondria to the nucleus in mammals. Future work will have to determine whether mTORC1-dependent ATF4 phosphorylation also contributes to the other pleiotropic functions of mTORC1 and ATF4^21,22,62,63^, under a variety of pathophysiological conditions.

## Methods

### Doxycycline treatment in C57BL/6J mice

Mice were housed under a 12-hour dark/12-hour light cycle and were allowed *ad libitum* to food and water. 9-10 weeks-old male C57BL/6J mice were randomly assigned to vehicle control or Doxycycline (Dox, Cat. D9891, Sigma) treatment group. The assigned mice (*n* = 5 per group) were administrated with either vehicle control or 50 mg/kg body weight of saline-dissolved Dox by intraperitoneal injection and sacrificed after 24 h of treatment. The experiment was carried out according to the institutional, Swiss national and European Union ethical guidelines and was approved by the local animal experimentation committee of the Canton de Vaud (License number, VD3478).

### RNA extraction and RNA-seq analysis

Cells or tissue powders were directly dissolved in 1 ml of the TriPure Isolation Reagent (Cat. 11667165001, Roche) and extracted using a column-based kit (Cat. 740955.250, Macherey-Nagel). RNA-seq was performed by BGI with the BGISEQ-500 platform.

For RNA-seq results, the raw data were filtered by removing adaptor sequences, contamination and low-quality (phred quality < 20) reads. Qualified reads were then mapped to the “*Mus_musculus*.GRCm38.95” genome with STAR aligner version 2.6.0a. Reads were counted using htseq-count version 0.10.0 using these flags: -f bam -r pos -s no -m union -t exon -i gene_id. Differential expression of genes was calculated by Limma-Voom. The genes with a Benjamini-Hochberg adjusted *P* value < 0.05 were defined as statistically significant. Genes whose expressions were significantly up-regulated (adjusted *P* value < 0.05) in Dox/AntiA/FCCP treatment condition; and were then down-regulated by more than 25% of the log_2_FC after ConA co-treatment, compared to the log_2_FC of Dox/AntiA/FCCP condition, were considered as v-ATPase activity-dependent. Functional clustering was conducted using the DAVID (Database for Annotation, Visualization and Integrated Discovery) database^68^. Heat-maps were generated using Morpheus (https://software.broadinstitute.org/morpheus).

### Quantitative RT-PCR

Total RNA was extracted as described above. cDNA was synthesized using the Reverse Transcription Kit (Cat. 205314, Qiagen). qRT-PCR was conducted with the LightCycler 480 SYBR Green I Master kit (Cat. 04887352001, Roche). Primers used for qRT-PCR are listed in Supplementary Table 2. Primers for mouse *Gapdh* and *Actin*, and human *GAPDH* and *ACTIN* were used as normalization controls.

### Cell culture and drug treatment

HEK293T cells (Cat. CRL-3216) were obtained from ATCC. Immortalized wild-type and *Atf4^−/−^* mouse embryonic fibroblasts (MEFs) were kindly provided by Prof. D. Ron (Cambridge Institute for Medical Research)^57^. All cell lines were validated to be free of mycoplasma contamination and maintained in Dulbecco’s modified Eagle’s medium containing 4.5 g glucose per liter and 10% fetal bovine serum. For culturing the *Atf4^−/−^* MEFs, 1× non-essential amino acids (Cat. 11140050, Gibco) and 55 μM β-mercaptoethanol (Cat. 31350010, Gibco) were furthermore supplemented to the medium, as described previously^57^; wild-type MEFs were cultured at the same condition for at least one week before comparing with the *Atf4^−/−^* MEFs. Plasmids expressing ATFS-1 and ATF4 were constructed by PCR amplifying from total cDNA of *C. elegans* and MEFs, respectively, and verified by sequencing. ATF4 mutants were created with the GeneArt™ Site-Directed Mutagenesis System (Cat. A13282, ThermoFisher). Plasmids expressing Myc-tagged Rheb (Plasmid #24941), Flag-tagged mTOR (Plasmid #26603) and HA-tagged Raptor (Plasmid #8513) were purchased from Addgene. Transfection were performed with the TransIT-X2 Transfection Reagent (Cat. MIR-6000, Mirus Bio). For CRISPR/Cas9-based knockdown of v-ATPase subunits, sgRNA for human *ATP6V0C* (5’-GAATAGTCGGGGCTGCTGGG-3’) and *ATP6V0D1* (5’-TCGATGACTGACACCGTCAG-3’) were cloned to lentiCRISPR v2 plasmid (Plasmid #52961, Addgene), followed by virus package and infection procedures as described previously^69^. The compounds used for treatment of cells were: Doxycycline (Cat. D9891, Sigma), Concanamycin A (Cat. C9705, Sigma), Bafilomycin A1 (Cat. S1413, Selleckchem), Cycloheximide (Cat. S7418, Selleckchem), Torin1 (Cat. S2827, Selleckchem), Rapamycin (Cat. S1039, Selleckchem), Antimycin A (Cat. A8674, Sigma), Oligomycin (Cat. 75351, Sigma), Tunicamycin (Cat. S7894, Selleckchem) and Chloroquine diphosphate (Cat. S4157, Selleckchem), with the concentrations indicated in the Figure legends.

### ChIP-qPCR of MEFs

ChIP-qPCR were performed as described previously^17^. Briefly, MEFs were fixed with 1% formaldehyde for 15 min and quenched by 0.125 mM glycine. Immunoprecipitations were carried out using antibody against ATF4 (1:100, Cat. 11815, CST). Sonication was conducted for a total time of 15 min. The primers used for ChIP-qPCR are listed in Supplementary Table 2.

### Western blots

Proteins were extracted with Radio-immunoprecipitation Assay (RIPA) buffer supplied with protease and phosphatase inhibitors, as described previously^29^. Immunoprecipitation of Flag-tagged proteins were carried out with the anti-FLAG M2 beads (Cat. A2220, Sigma) in RIPA buffer. For western blotting, the antibodies used were: P-EIF2α (Cat. 3597, CST, 1:500), Tubulin (Cat. T5168, Sigma, 1:2,000), P-S6K (Cat. 9205, CST, 1:1,000), S6K (Cat. 9202, CST, 1:1,000), P-S6 (Cat. 2215, CST, 1:1,000), S6 (Cat. 2317, CST,1:1,000), P-4E-BP1 (Cat. 9644, CST, 1:1,000), 4E-BP1 (Cat. 2855, CST, 1:1,000), ASNS (Santa Cruz, Cat. sc-365809, 1:1,000), EIF2α (Cat. 9722, CST, 1:1,000), ATF4 (Cat. 11815, CST, 1:1,000), ATF5 (Cat. ab60126, Abcam, 1:1,000), ATP6V0D1 (Cat. ab202899, Abcam, 1:1,000), Flag-tag (F7425, Sigma, 1:1,000), Myc-tag (Cat. sc-40, Santa Cruz, 1:2,000), mTOR (Cat. 2972, CST, 1:1,000), Phospho-S*P (Cat. 2325, CST, 1:1,000; for detecting P-S166-ATF4), Phospho-ST*P (Cat. 5243, CST, 1:1,000; for detecting P-T173-ATF4), and HRP-labelled anti-rabbit (Cat. 7074, CST, 1:5,000), anti-rabbit (Light-Chain Specific) (Cat. 93702, CST, 1:5,000, for detecting the endogenously immunoprecipitated ATF4 and its phosphorylation) and anti-mouse (Cat. 7076, CST, 1:5,000) secondary antibodies.

### Imaging of mammalian cells

For Mitotracker staining of MEFs, cells were grown on glass cover slips and MitoTracker Red CMXRos (Cat. M7512, Invitrogen) was added to the culture medium 30 min prior to imaging according to manufacturer’s instructions. Cells were then fixed and stained with antibodies to early endosome marker Rab5 (Cat. 3547, CST, 1:200), late endosome marker Rab7 (Cat. 9367, CST, 1:200), or lysosome marker Lamp1 (Cat. 121617, Biolegend, 1:250). For imaging the lysosomal-localized mTOR, fixed MEFs were stained with mTOR (Cat. 2972, CST, 1:200), early endosome marker Rab5 (Cat. 46449, CST, 1:200), late endosome marker Rab7 (Cat. 95746, CST, 1:200), or lysosome marker Lamp1 (Cat. 121617, Biolegend, 1:250) antibody. Images were then acquired using a ZEISS LSM 700 confocal microscope and analyzed by using ImageJ with a Mito-Morphology macro^70^. For mitochondrial network analysis, at least 20 cells were analyzed for each condition.

### Mitochondrial respiration assay

Oxygen consumption rate (OCR) of cultured MEFs was determined using the Seahorse XFe96 Extracellular Flux Analyzer (Agilent Technology) according to the manufacturer’s protocol. The OCR was measured upon serial injections of 2 μM Oligomycin, 2 μM FCCP, and a mixture of 1 μM Rotenone/Antimycin A. The OCR values were normalized to total cell number. For the measurement of OCR in response to Antimycin A treatment, MEFs were treated with 2 μM Antimycin A for 24 h, and then washed three times with control medium. Cells were then let recovered in control medium for 8 h and a standard OCR measurement assay was subsequently conducted.

### Mitochondrial ROS quantification

Mitochondrial ROS levels were measured using MitoSox (Cat. M36008, Thermofisher). Cells were treated with 2 μM Antimycin A for 48 h. MEFs were then trypsinized and incubated with 2 μM MitoSox at 37 °C for 30 min. After washed twice with PBS, the cells were then analyzed with flow cytometry. Data were quantified and plotted with FlowJo software. Three independent experiments were conducted and similar results were acquired.

### *In vitro* kinase assay

Kinase assays were performed as described previously^71^, with slight modifications. For kinase assay using mTORC1, HEK293T cells were transfected with plasmids expressing Flag-tagged mTOR (Plasmid #26603, Addgene) together with HA-tagged Raptor (Plasmid #8513, Addgene). 36 h post-transfection, cells were lysed in CHAPS lysis buffer [40 mM HEPES (pH 7.5), 0.3% CHAPS, 120 mM NaCl, 1 mM EDTA, 10 mM pyrophosphate, 10 mM glycerophosphate, 50 mM NaF, 1.5 mM Na_3_VO_4_ and 1 × protease inhibitor Cocktail (Cat. 78430, ThermoFisher)]. mTORC1 were then immunoprecitated using anti-Flag M2 Beads (Cat. A2220, Sigma). The immunoprecipitates were washed 3 times with the CHAPS lysis buffer and the mTOR reaction buffer [25 mM HEPES (pH 7.4), 50 mM KCl, 20% glycerol, 10 mM MgCl_2_, 4 mM MnCl_2_, 1 mM DTT], respectively. The assays were carried out in 50 μl mTOR reaction buffer with or without 200 μM ATP and/or the recombinant human His-tagged ATF4 (Cat. ab109946, Abcam) at 30°C for 45 min. When indicated, Torin1 (250 nM, Cat. S2827, Selleckchem) was added 10 min prior to the start of the assay. Reactions were stopped by adding 4 × SDS loading buffer. For kinase assays using recombinant mTOR, the recombinant GST-tagged kinase active human mTOR (Cat. PV4753, ThermoFisher) was used instead of the immunoprecitated mTORC1. Following SDS-PAGE and SimplyBule SafeStain (Cat. LC6060, Invitrogen) staining, the bands corresponding to His-ATF4 were sliced, digested with either trypsin or GluC, and then analyzed by Liquid Chromatograph Triple Quadrupole Mass Spectrometer (LC-MS/MS) for phosphorylated peptides.

### Statistical analysis

All statistical analyses were performed using Graphpad Prism 8 software. Differences between two groups were assessed using two-tailed unpaired Student’s *t*-tests. No statistical methods were used to predetermine sample size. The experiments were not randomized, and investigators were not blinded to allocation during experiments and outcome assessment. Data distribution was assumed to be normal but this was not formally tested. Analysis of variance (ANOVA) followed by Tukey post-hoc test (one-way ANOVA for comparisons between groups, and two-way ANOVA for comparisons of magnitude of changes between different groups from different treatments or cell lines) was used when comparing more than two groups.

### Reporting summary

Further information on research design is available in the Nature Research Reporting Summary linked to this article.

## Data availability

Original reagents are available upon request. The raw and processed sequencing datasets have been deposited in the NCBI Gene Expression Omnibus (GEO) database with the accession numbers: GSE179510 (Reviewer access: udutaqywhhghhgr).

## Acknowledgments

We thank D. Ron (Cambridge Institute for Medical Research) for providing the *Atf4^−/−^* MEFs. We thank all members of J. Auwerx and K. Schoonjans laboratories for helpful discussions. This work was supported by grants from the Ecole Polytechnique Federale de Lausanne (EPFL), the European Research Council (ERC-AdG-787702), the Swiss National Science Foundation (SNSF 31003A_179435) and GRL grant of the National Research Foundation of Korea (NRF 2017K1A1A2013124). T.Y.L. was supported by the “Human Frontier Science Program” (LT000731/2018-L). Q.W. was supported by the European Molecular Biology Organization (ALTF 111-2021). A.W.G. was supported by the Accelerator prize given by the United Mitochondrial Disease Foundation (PF-19-0232). X.L. was supported by the China Scholarship Council (201906050019).

## Author contributions

T.Y.L. and J.A. conceived the project. T.Y.L., Q.W. and A.W.G. performed most of the experiments. X.L. performed the RNA-seq data analysis. A.M. contributed to the mouse *in vivo* study. M.S., and J.A. supervised the study. T.Y.L. and J.A. wrote the manuscript with comments from all authors.

---

**Extended Data Fig. 1.**
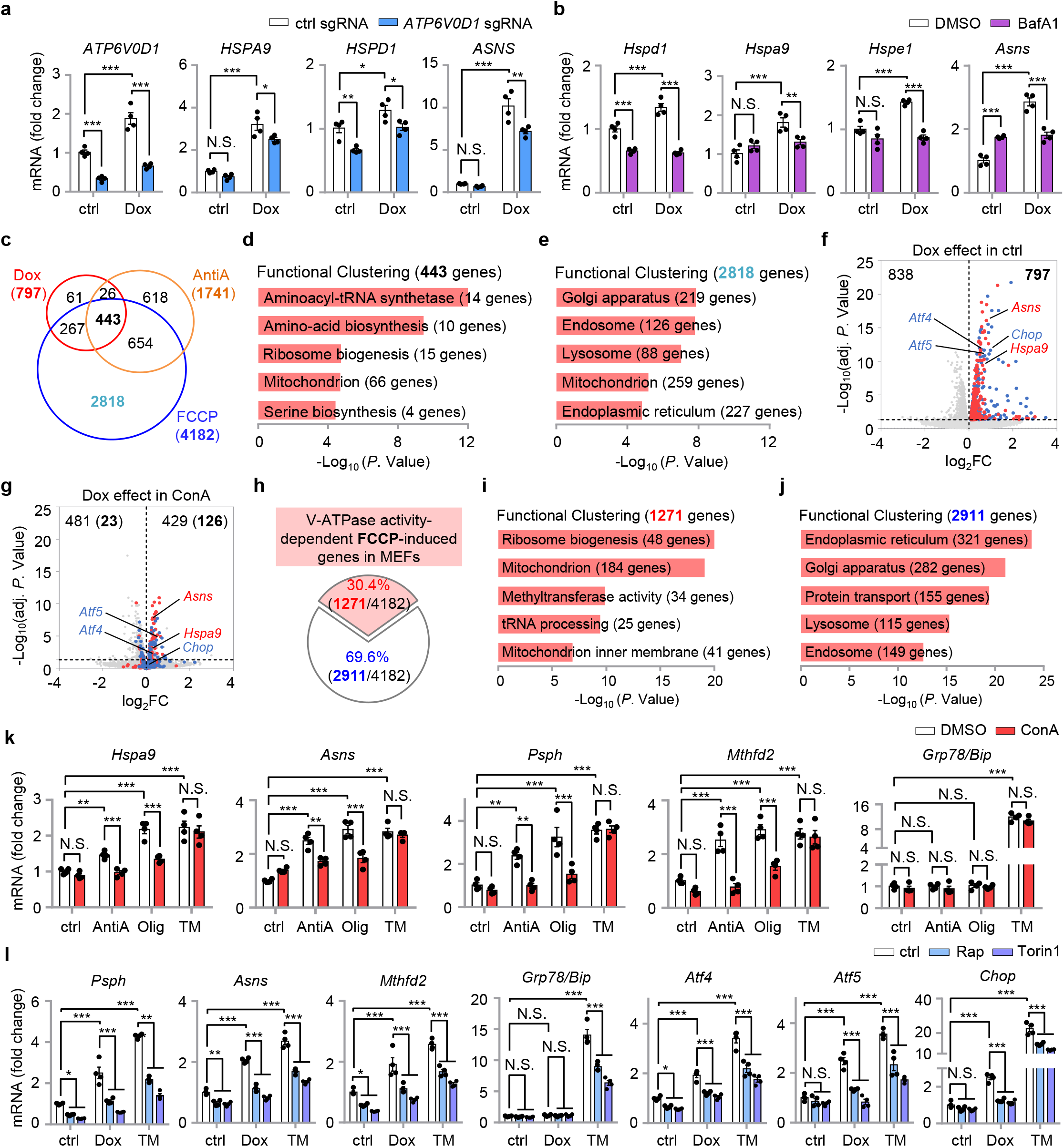
An essential role of v-ATPase and mTORC1 activity in UPR^mt^ activation. **a**, qRT-PCR results (*n* = 4 biologically independent samples) of HEK293T cells expressing control (ctrl) or *ATP6V0D1* sgRNA, and treated with or without Doxycycline (Dox) (30 μg/ml) for 24 h. **b**, qRT-PCR results (*n* = 4 biologically independent samples) of MEFs pretreated with DMSO control or Bafilomycin A1 (BafA1) (1 μM) for 1 h, and then co-treated with or without Dox (30 μg/ml) for 24 h. **c**, The up-regulated transcripts in response to Dox, Antimycin A (AntiA) or FCCP treatment based on the RNA-seq dataset. **d**,**e**, Functional clustering of the 443 (**d**) and 2,818 (**e**) genes as indicated in (**c**). **f**,**g**, Volcano plots showing the effect of Dox in control (DMSO-treated) MEFs (**f**), or in ConA-treated MEFs (**g**) (numbers in bold are the common genes within the 797 genes as indicated in (**f**)). FC, fold change. Genes whose induction upon Dox treatment is dependent on v-ATPase activity are highlighted in red, the three putative UPR^mt^ transcription factors whose induction is independent on v-ATPase activity are highlighted in blue. **h**, Diagram of the UPR^mt^ genes that are dependent (red) or independent (blue) on v-ATPase activity for induction upon FCCP treatment. **i**,**j**, Functional clustering of the 1,271 (**i**) and 2,911 (**j**) genes as indicated in (**h**). **k**, qRT-PCR results (*n* = 4 biologically independent samples) of MEFs pretreated with DMSO control or ConA (200 nM) for 1 h, and then co-treated with or without AntiA (2 μM), Oligomycin (Olig, 2 μM), or Tunicamycin (TM, 1.5 μg/ml) for 24 h. **l**, qRT-PCR results (*n* = 4 biologically independent samples) of MEFs pretreated with DMSO, Rapamycin (Rap, 100 nM) or Torin1 (250 nM) for 1 h, and then co-treated with or without Dox (30 μg/ml) or TM (1.5 μg/ml) for 24 h. Error bars denote S.E.M. Statistical analysis was performed by ANOVA followed by Tukey post-hoc test (\**P* < 0.05; \*\**P* < 0.01; \*\*\**P* < 0.001; N.S., not significant).

**Extended Data Fig. 2.**
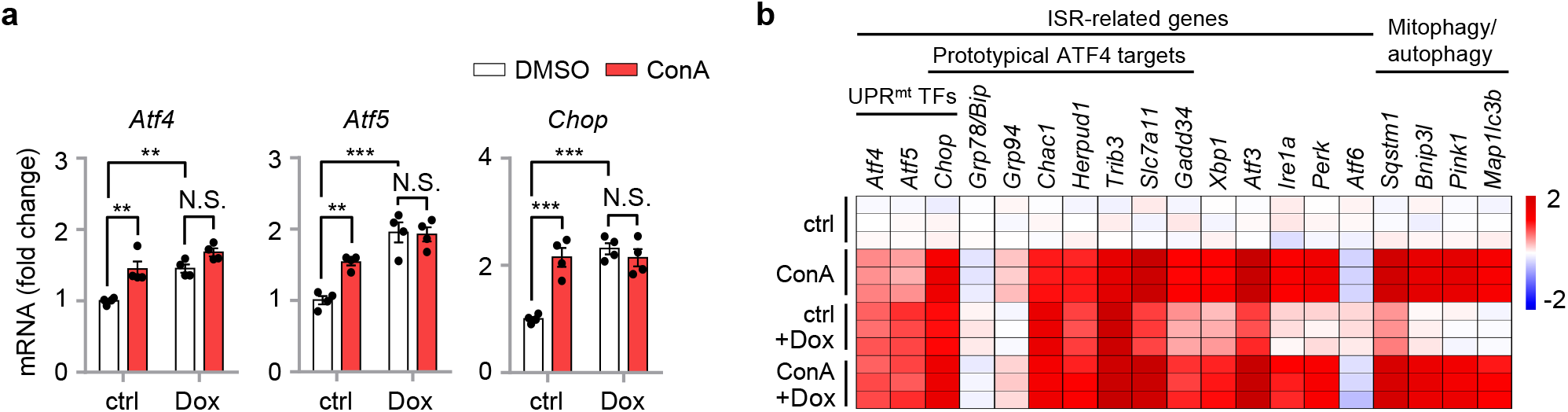
Impacts of Dox and v-ATPase inhibitor ConA in gene expression. **a**, qRT-PCR results (*n* = 4 biologically independent samples) of MEFs pretreated with DMSO control or v-ATPase inhibitor Concanamycin A (ConA) (200 nM) for 1 h, and then co-treated with or without Dox (30 μg/ml) for 24 h. **b**, Heat-map of the relative expression levels of genes in MEFs treated with Dox and/or ConA in log_2_ fold-change, based on the RNA-seq dataset. ISR, integrated stress response. See Supplementary Table 1 for detailed gene expression changes.

**Extended Data Fig. 3.**
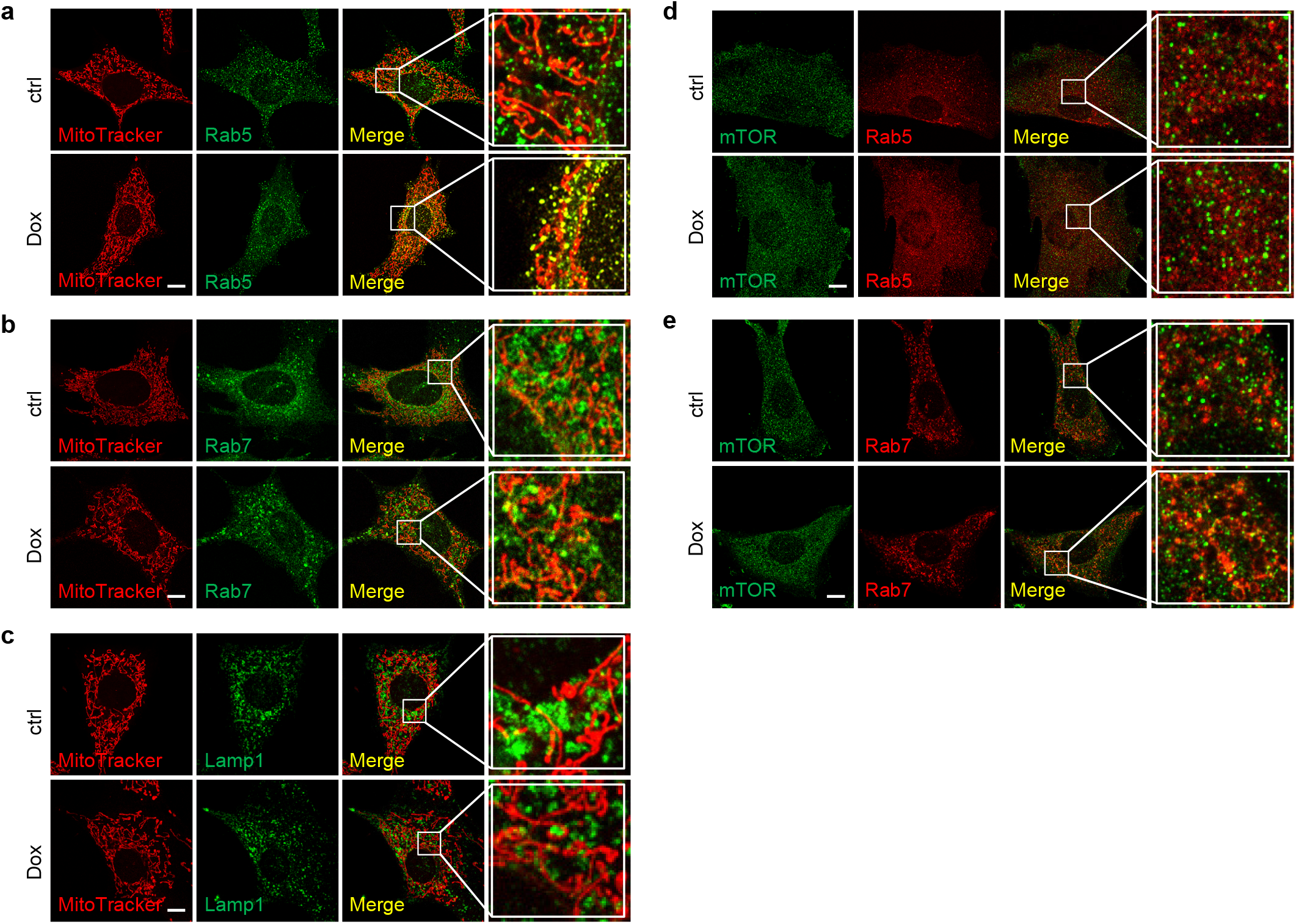
A mitochondrion-endosome-lysosome route for mTORC1 activation upon mitochondrial stress. **a**-**c**, MitoTracker co-localizes with early endosomes (**a**), partially co-localizes with late endosomes (**b**), and does not co-localize with lysosomes (**c**), during mitochondrial stress. MEFs were stained with MitoTracker (red) for 1 h, and then treated with or without Dox (30 μg/ml) for 3 h, cells were then fixed and stained with the early endosome marker Rab5 (green) (**a**), the late endosome marker Rab7 (green) (**b**), or the lysosome marker Lamp1 (green) antibodies (**c**). **d**,**e**, mTOR does not co-localize with the early endosomes (**d**) and partially co-localizes with late endosomes (**e**) during mitochondrial stress. MEFs were treated with or without Dox (30 μg/ml) for 3 h, cells were then fixed and co-stained with mTOR (green), and early endosome marker Rab5 (red) (**d**), or late endosome marker Rab7 (red) antibodies (**e**). Scale bars, 10 μm.

**Extended Data Fig. 4.**
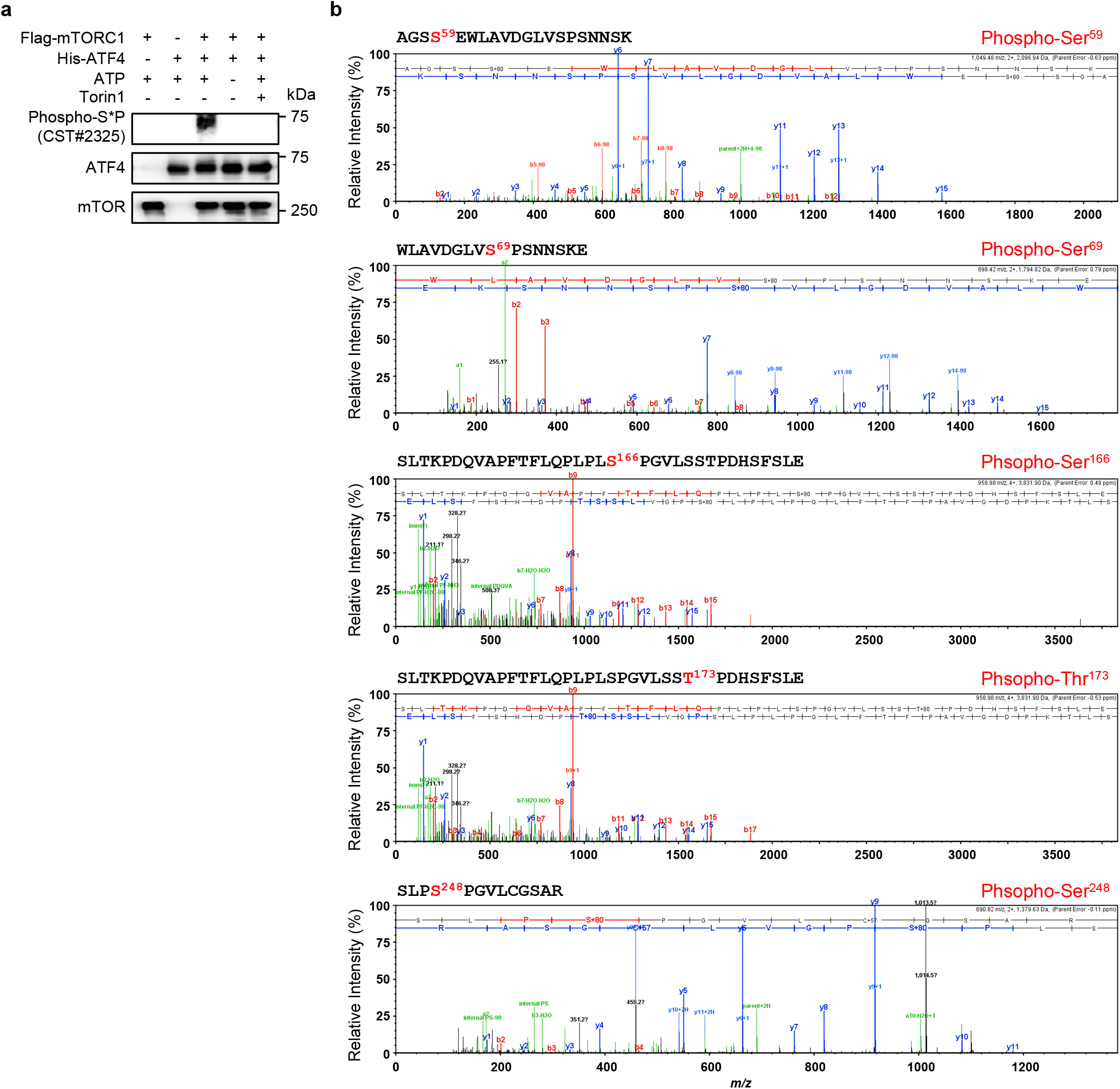
Identification of ATF4 as a direct phosphorylation substrate of mTORC1. **a**, mTORC1 directly phosphorylates ATF4 *in vitro*. *In vitro* kinase assay was performed with Flag-tagged mTORC1 immunoprecipitated from HEK293T cells and recombinant His-tagged ATF4, with or without Torin1 (250 nM). **b**, The representative spectrums for the phosphorylated peptides of human ATF4 identified by Liquid Chromatograph Triple Quadrupole Mass Spectrometer (LC-MS/MS), with numbering according to the amino acid sequence of human ATF4 protein

**Extended Data Fig. 5.**
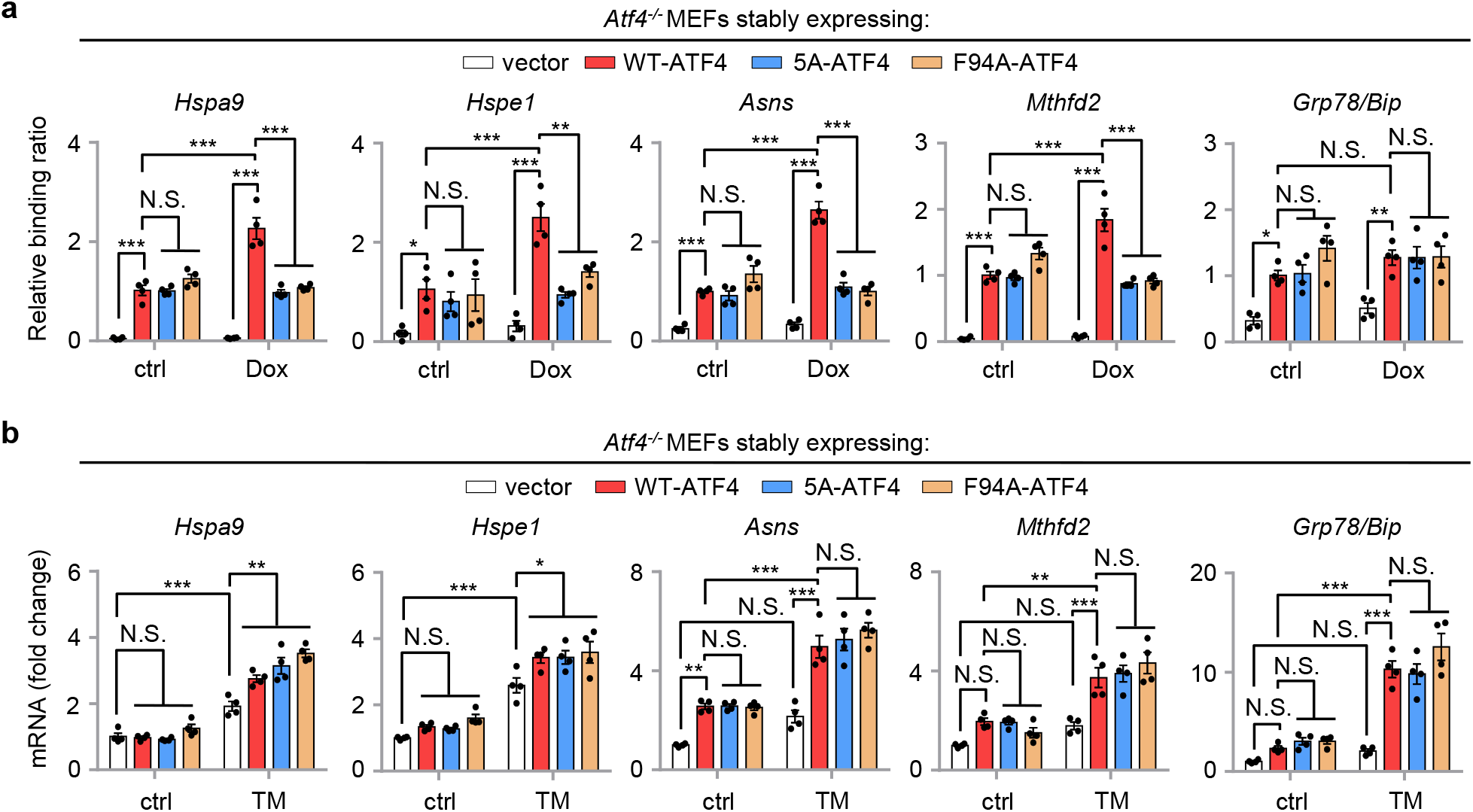
mTORC1-mediated ATF4 phosphorylation is required for mitochondrial stress-induced ATF4 binding to the promoters of UPR^mt^ genes, but non-essential for ER stress response. **a**, ATF4 phosphorylation defective (5A) and TOS motif disrupted (F94A) mutants failed to bind to the promoters of UPR^mt^ genes in *Atf4^−/−^* MEFs upon Dox treatment. ATF4 ChIP-qPCR analysis (*n* = 4 biologically independent samples) of the promoters of the UPR^mt^ genes in *Atf4^−/−^* MEFs stably expressing empty vector, wild-type, 5A or F94A forms of ATF4, with or without Dox (30 μg/ml) treatment for 3 h. **b**, qRT-PCR results (*n* = 4 biologically independent samples) of *Atf4^−/−^* MEFs stably expressing empty vector, wild-type, 5A or F94A forms of ATF4, with or without Tunicamycin (TM, 1.5 μg/ml) treatment for 24 h. Error bars denote S.E.M. Statistical analysis was performed by ANOVA followed by Tukey post-hoc test (\**P* < 0.05; \*\**P* < 0.01; \*\*\**P* < 0.001; N.S., not significant).

**Extended Data Fig. 6.**
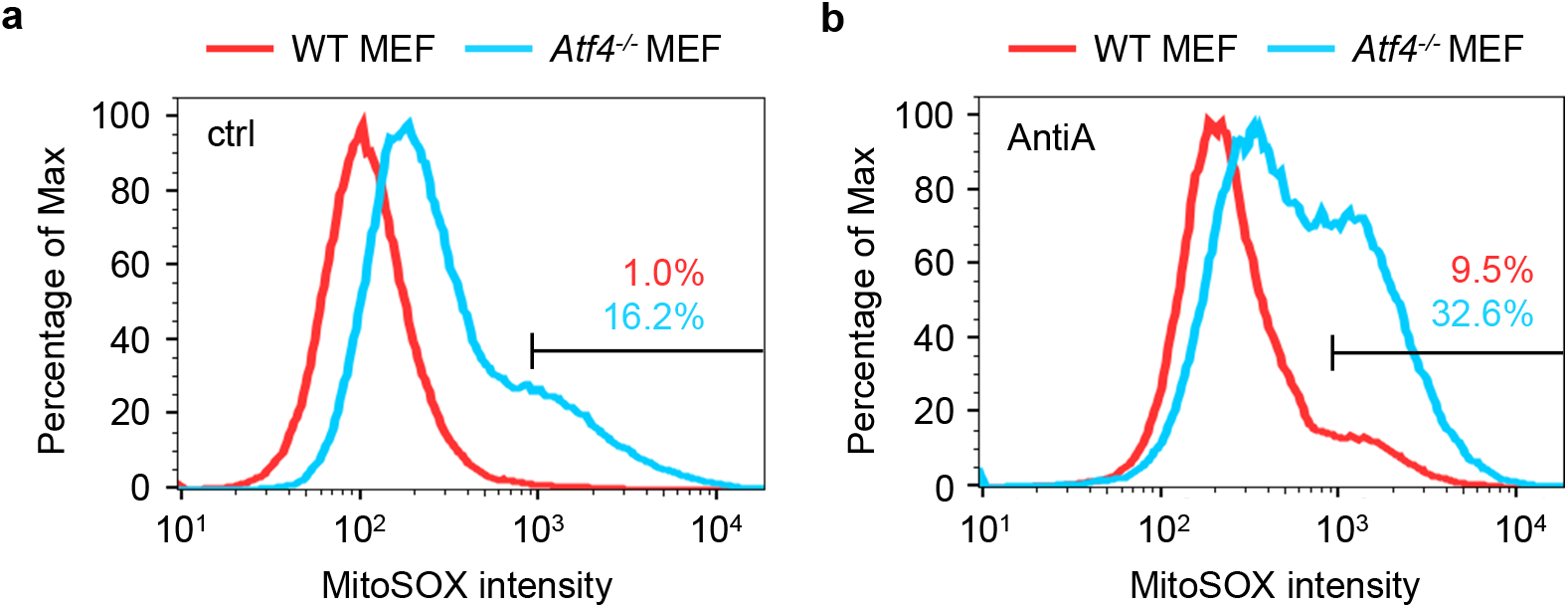
*Atf4* loss-of-function leads to disruption of mitochondrial redox homeostasis. **a**,**b**, Representative flow cytometry result of the mitochondrial superoxide (MitoSOX) intensity of wild-type (WT) and *Atf4^−/−^* MEFs, after DMSO control (ctrl) (**a**) or Antimycin A (AntiA, 2 μM) (**b**) treatment for 48 h. The percentages of MitoSOX-positive cells are indicated.

**Supplementary Table 1. RNA-seq results for MEFs treated with or without mitochondrial stress inducers and/or ConA**.

**Supplementary Table 2. List of primers used for qRT-PCR and ChIP-qPCR in this study**.

