## Supplemental Table 2 for "Lysosomes mediate the mitochondrial UPR via mTORC1-dependent ATF4 phosphorylation"

**Supplementary Table 2. List of primers used for qRT-PCR and ChIP-qPCR in this study.**

| Species | Application | Gene | Forward primer (5'-3') | Reverse primer (5'-3') |
|---|---|---|---|---|
| mouse | qRT-PCR | *Hspd1* | TCTTCAGGTTGTGGCAGTCA | CCCCTCTTCTCCAAACACTG |
| mouse | qRT-PCR | *Hspa9* | AATGAGAGCGCTCCTTGCTG | CTGTTCCCCAGTGCCAGAAC |
| mouse | qRT-PCR | *Hspe1* | CTGACAGGTTCAATCTCTCCAC | AGGTGGCATTATGCTTCCAG |
| mouse | qRT-PCR | *Asns* | TTGACCCGCTGTTTGGAATG | CGCCTTGTGGTTGTAGATTTCAC |
| mouse | qRT-PCR | *Mthfd2* | CCTACAGCCCTTCCACCTG | TCCTGCTGTACTTCTTGCTTGA |
| mouse | qRT-PCR | *Psph* | GAGGCCGCAGTGTCTGAAAT | CACAATGCTCCGAAAGCCAC |
| mouse | qRT-PCR | *Cth* | ATAGTCGGCTTCGTTTCCTG | TCGGCAGCAGAGGTAACAAT |
| mouse | qRT-PCR | *Grp78/Bip* | ACTTGGGGACCACCTATTCCT | ATCGCCAATCAGACGCTCC |
| mouse | qRT-PCR | *Atf4* | GAAACCTCATGGGTTCTCCA | GAAAAGGCATCCTCCTTGC |
| mouse | qRT-PCR | *Atf5* | AGAGCCCCTGGCAGGTGA | CAGAGGAAGGAGAGCTGTGAAGT |
| mouse | qRT-PCR | *Chop* | CGGAACCTGAGGAGAGAGTG | CGTTTCCTGGGGATGAGATA |
| mouse | qRT-PCR | *Gapdh* | TGTGTCCGTCGTGGATCTGA | CCTGCTTCACCACCTTCTTGAT |
| mouse | qRT-PCR | *Actin* | GAGACCTTCAACACCCC | GTGGTGGTGAAGCTGTAGCC |
| human | qRT-PCR | *ATP6V0C* | GCTTCGTTTTTCGCCGTCAT | GTCATTCAGGGAGTTGGCGA |
| human | qRT-PCR | *ATP6V0D1* | CTACCTCAACCTGGTGCAGT | GTTCCTCATGTGGCGGAACT |
| human | qRT-PCR | *HSPA9* | TGGTGAGCGACTTGTTGGAAT | ATTGGAGGCACGGACAATTTT |
| human | qRT-PCR | *HSPD1* | GGGTAACCGAAGCATTTCTGC | CTGCACTCTGTCCCTCACTC |
| human | qRT-PCR | *ASNS* | ATCACTGTCGGGATGTACCC | TGATAAAAGGCAGCCAATCC |
| human | qRT-PCR | *GAPDH* | TTGGTATCGTGGAAGGACTC | ACAGTCTTCTGGGTGGCAGT |
| human | qRT-PCR | *ACTIN* | GTCATCACCATTGGCAATGAG | CGTCATACTCCTGCTTGCTG |
| mouse | ChIP-qPCR | *Hspa9* | GCTTCACGACCTCTGTCCG | AAAGACTCAAGGTCACACGGG |
| mouse | ChIP-qPCR | *Hspe1* | GCTCCCCTTTTCTTCCGCCT | GCTCCGGACTCTGAACTCGG |
| mouse | ChIP-qPCR | *Asns* | CAGAACACCTCCTGGCTCTC | AGTGACAAGACCGGTTGGAG |
| mouse | ChIP-qPCR | *Mthfd2* | CTGCCACTGCAGAGATGGGTG | GAGGGAAGTTGGTACCCTTGGAG |
| mouse | ChIP-qPCR | *Grp78/Bip* | GCTCGATACTGGCCGAGACA | CGACGACGGTTCTGGTCTG |
| mouse | ChIP-qPCR | *Trib3* | GGTCACAGATGGTGCAATCC | CTCTTCCGCTGCTAAAGTGC |
| mouse | ChIP-qPCR | *Slc7a11* | GCTGAGTAATGTTGGCGCTTTCTC | CACACCAACTTACTGGGCTGC |
